# Community and functional stability in a working bioreactor degrading 1,4-dioxane at the Lowry Landfill Superfund Site

**DOI:** 10.1101/2025.03.20.644286

**Authors:** Jessica L. Romero, Jack H. Ratliff, Christopher J. Carlson, Daniel R. Griffiths, Christopher S. Miller, Annika C. Mosier, Timberley M. Roane

## Abstract

1,4-dioxane (dioxane) is an emerging contaminant that poses risks to human and environmental health. Bacterial dioxane degradation is increasingly being studied as a method to remove dioxane from contaminated water. However, there is a lack of studies on microbial community structures and functions within efficient, large scale, biodegradation-based remediation technologies. The Lowry Landfill Superfund Site (Colorado, USA) uses an on-site, pump-and-treat facility to remove dioxane from contaminated groundwater by biodegradation. Here, 16S rRNA and shotgun metagenomic sequencing were used to describe microbial community composition, soluble di-iron monooxygenase (SDIMO) alpha hydroxylases, and potential for dioxane degradation and horizontal gene transfer in bioreactor support media from the facility. Support media showed diverse microbial communities dominated by Nitrospiraceae, Nitrososphaeraceae, and Nitrosomonadaceae. *Pseudonocardia* were also detected, suggesting a potential presence of known dioxane-degraders. Candidate SDIMOs belonged mostly to Group V, followed by Groups IV, II, and I (based on read depth). The most abundant Group V clade contained 38 proteins that were phylogenetically related to DxmA-like proteins, including that of *Pseudonocardia dioxanivorans* CB1190 (a known dioxane degrader). Seventeen Lowry contigs containing DxmA-like proteins contained protein-coding genes potentially involved in chemical degradation, transcriptional regulation, and chemical transport. Interestingly, these contigs also contained evidence of potential horizontal gene transfer, including toxin-antitoxin proteins, phage integrase proteins, putative transposases, and putative miniature inverted-repeat transposable elements. These findings improve our understanding of potential dioxane biodegradation mechanisms in a functioning remediation system. Further studies are needed to definitively confirm microbial activity and enzymatic activity towards dioxane removal in this site.

**IMPORTANCE:** As an environmental contaminant, 1,4-dioxane poses risks for water quality and human health. Used as a solvent and chemical stabilizer in a variety of manufacturing and industrial applications, microbiological methods of detoxification and mitigation are of interest. The degradation of 1,4-dioxane by the bacterium *Pseudonocardia spp.* is the best understood example; however, these studies are largely based on single isolate, bench-scale, or *in silico* experiments. Consequently, a knowledge gap exists on bacterial degradation of 1,4-dioxane at environmentally relevant concentrations using functioning remediation technologies at scale. This study addresses this gap directly by describing microbial taxa, enzymes, and potential horizontal gene transfer mechanisms associated with an active treatment plant located on a 1,4-dioxane-impacted U.S. Environmental Protection Agency (EPA) superfund site. As 1,4-dioxane contamination gains more attention, these findings may prove useful for future facilities aiming to promote and optimize removal by biodegradation.

## INTRODUCTION

The organic chemical 1,4-dioxane (dioxane) is an emerging contaminant that has recently gained much attention due to its environmental prevalence and recognition as a possible carcinogen (1). Used since the 1920s as a degreaser, then shortly as a chlorinated solvent stabilizer, dioxane is used in many industrial processes including the production of plastics, glues, cosmetics, antifreeze, pharmaceuticals, textiles, pesticides, and other materials (2–4). Given the various uses of dioxane, it is frequently released from industrial and consumer activities. For instance, U.S. industries were estimated to have released 705,000 pounds of dioxane into the environment in 2015 (5). However, this is likely an underestimate as it does not include, for example, releases as an unwanted chemical byproduct in industrial wastewater (5, 6). In addition, consumer use of products frequently created or contaminated with dioxane during manufacturing, such as soaps and laundry detergents, releases dioxane into waterways and landfills (2, 7). Waterway releases and landfill leachate may enter municipal wastewater treatment systems (7), which are estimated to remove 5% of dioxane inputs at most (5). Dioxane is not readily broken down by these treatment plants due to its miscibility and the stability of its two ether bonds (8). Due in part to these properties, plus its hydrophilic nature and low volatility, dioxane tends to associate with water (9). Consequently, dioxane has been recognized as a drinking water contaminant as early as 1978 (5). In 2015, the U.S. Environmental Protection Agency (EPA) detected dioxane in 21% of tested drinking water sources in the U.S. (3). Although EPA classifies dioxane as a potential human carcinogen, the agency does not currently enforce a federal dioxane release limit (3, 7). Therefore, dioxane limits for surface and groundwater differ across the U.S. (1, 3). In 2004, Colorado became the first state to set a dioxane limit of up to 6.1 µg/L for surface water and groundwater to be met by March 2005 (2). The Colorado dioxane limit has since decreased to 0.35 µg/L (1). Recently, states like New York have begun to restrict dioxane levels in personal care products and cleaning products (7, 10).

Several methods have been tested to remove dioxane from contaminated water with varying successes. Certain physical removal methods are inefficient as they only change the physical phase of or concentrate dioxane without degrading it (3). Chemical methods known as advanced oxidation processes (AOPs) completely degrade dioxane to carbon dioxide with high efficiency but are relatively expensive and less effective in treating chemical mixtures (3, 5, 8, 11, 12). Lastly, biological methods remove dioxane through degradation by microorganisms (biodegradation). Two biodegradation mechanisms exist: direct metabolism, a growth-linked process in which dioxane is directly degraded to yield carbon and energy (13), and co-metabolism, a non-growth-linked process in which dioxane is partially degraded in a fortuitous manner (13, 14). Co-metabolism occurs in the presence of a non-dioxane co-substrate that induces dioxane degradation (13). Co-substrates that have induced dioxane degradation include tetrahydrofuran (THF), methane, propane, butanol, and toluene (15).

To date, approximately 30 bacterial species and three fungal species have shown dioxane degradation (13, 15, 16). The best characterized dioxane-degrading bacterial strain is *Pseudonocardia dioxanivorans* CB1190 (CB1190), isolated from dioxane-contaminated industrial sludge (17). Under aerobic conditions, CB1190 directly metabolized 50% of dioxane in a 4.0 mM (~0.35 g/L) culture in 18 hours (17). An example of a dioxane co-metabolizing bacterium is *Methylosinus trichosporium* OB3b (OB3b), which co-metabolized 50 mg/L (~0.57 mM) of dioxane at a rate of 0.38 ± 0.02 mg/h/mg protein when supplied with a 25% (vol/vol) methane gas addition and no copper salt addition (18).

While dioxane degradation has been studied extensively in microbial isolate strains, recent work has also shown synergistic degradation by complex microbial communities (19). Using an activated sludge enrichment, one study proposed a dioxane degradation pathway in which the bacteria *Xanthobacter spp*. and *Rhizobiales* contributed enzymes that initiated hydroxylation, which was then followed by subsequent pathway steps carried out by other bacteria (19). Another study suggested that certain microbial community members, like *Ancylobacter polymorphus* ZM13, initiated dioxane degradation whereas other members, like *Xanthobacter* and *Mesorhizobium*, metabolized intermediates and alleviated stress responses (20).

Few studies have directly applied dioxane biodegradation towards wastewater and landfill contamination. One study achieved 81.26 ± 6.17% dioxane removal from polyester factory wastewater using a pilot-scale, multi-staged reactor inoculated with anaerobic digester sludge (21). Another study reduced dioxane below detection levels (<0.38 µg/L) in landfill groundwater using microcosms bioaugmented with bacterial dioxane-degrading strains (22). Despite these advances, there is currently a lack of literature on microbial community responses within large-scale remediation technologies showing stable, efficient dioxane biodegradation.

The enzymes that initiate microbial dioxane degradation are known as soluble di-iron monooxygenases (SDIMOs). Six SDIMO groups exist based on phylogeny, substrate range, sequence identity, and other traits (15, 23). To date, SDIMOs with potential dioxane degradation capabilities have been identified from all groups, except for Group IV (24). Recently, eight candidate dioxane-degrading SDIMOs were described and searched for broadly in environmental metagenomes (15). This and other studies have suggested that SDIMO composition in microbial communities may be affected by environmental conditions (15, 25–27). Further study of SDIMO presence, structures, and functions in the context of dioxane remediation could elucidate dioxane biodegradation mechanisms and potentially reveal strategies to enhance these.

Here, we evaluated the bacterial degradation of dioxane in an on-site flow-through wastewater bioremediation plant at the Lowry Landfill Superfund Site (Aurora, Colorado) (12). In this study, 16S rRNA and shotgun metagenomic sequencing were used to: 1) identify the microbial communities potentially involved in dioxane degradation at the Lowry Landfill treatment plant, 2) describe the Lowry SDIMOs potentially involved in dioxane degradation and compare these to SDIMOs from the literature, and 3) examine the Lowry protein-coding genes genomically located near the SDIMOs to characterize potential dioxane degradation mechanisms. This work is novel in that it describes the structures and functions of an in-compliance bioremediation treatment plant. These findings will improve our understanding and management of biological remediation of 1,4-dioxane.

## MATERIALS AND METHODS

### Site description

The Lowry Landfill Superfund Site in Aurora, Colorado, is a dioxane-impacted site that uses *in situ* microorganisms in the aboveground bioreactor treatment of dioxane-containing groundwater. Accepting industrial and municipal wastes from 1964 to 1980, wastes were disposed of in 200 acres of unlined pits. While these disposal practices followed regulatory standards of the time, they eventually led to chemical contamination of the surrounding soil, sediment, surface and groundwater (28, 29). In 1984, the U.S. Environmental Protection Agency designated the landfill a superfund site, prompting a series of remediation efforts (30). This included the installation of an on-site hydraulic control through groundwater extraction system and aboveground treatment that originally used air stripping and granular activated carbon (31).

While the original treatment processes successfully removed volatile organic compounds from contaminated groundwater, it was later found that treatment did not reduce dioxane. Furthermore, treated groundwater was re-injected in accordance with Colorado State groundwater policy until 2004 (31). During this time, it was also found that this practice resulted in a dioxane plume extending from the north end of the landfill.

To address these issues, a new aerobic (micro)biological treatment system (BTS) was installed, which was composed of three aerated moving bed bioreactors (bioreactors) (Fig. S1A, B) (28, 32). Bioreactor operation has been detailed previously, including the use of Kaldnes K1 polyethylene support media (support media) (Wigan, UK) (Fig. S1C) to provide a surface for microbial growth, coarse bubble diffusion for aeration, and controls for temperature at 23.5°C and pH at 7.0 (12). The three bioreactors concurrently treat groundwater collected from locations distributed across the landfill (12). The removal of dioxane, and its known co-substrate tetrahydrofuran (THF), are specifically monitored in accordance with Colorado State permitted standards (12). Quantification of both compounds was performed by Eurofins Scientific (Arvada, CO) (https://www.eurofins.com/) using gas chromatography and mass spectrometry following EPA Method 8260B SIM for dioxane and EPA Method 8260B for THF (33). Dioxane removal efficiency of the BTS generally ranges from 90-98% (12) and effluent measures below 25 µg/L of dioxane on average (31). Since 2004, the landfill has discharged treated effluent to a local municipal wastewater treatment plant under a discharge permit allowing a limit of 220 µg/L of dioxane (12).

### Sample collection

To investigate dioxane biodegradation in the Lowry Landfill BTS, support media were collected from one of the three bioreactors (Bioreactor 1) (Fig. S1). Samples were collected at four timepoints over three years: 2019-03-26, 2022-01-18, 2022-01-25, and 2022-03-22. Although these timepoints do not represent a true time series, they were selected to allow preliminary comparisons of microbial communities across short and long timescales. At each sampling, support media were removed from Bioreactor 1 and placed into sterile 50 mL centrifuge tubes. Excess water was decanted from centrifuge tubes before sealing. Collected samples were stored at −20°C.

### 16S rRNA gene sequencing

To describe the microbial community composition of the Lowry Landfill treatment plant BTS, high-throughput sequencing of the 16S rRNA gene was performed on support media. As part of a CURE (course-based undergraduate research experience) during the Fall 2022 semester, students in the University of Colorado Denver General Biology teaching laboratory extracted total DNA for 16S rRNA gene amplification and sequencing. This allowed for a larger number of replicates sampled from each day. Using a quarter section of a single support media cut with a sterilized razorblade, DNA was extracted following the protocol from the QIAGEN DNeasy PowerSoil Kit (Hilden, Germany) with two modifications: 1) samples were incubated at 60°C for 10 minutes prior to bead beating for 10 minutes on a laboratory vortex at maximum speed with an 24-tube adapter, and 2) DNA was eluted in 50 µL of kit elution buffer (10 mM Tris). To distinguish support media quarter sections from multiple replicates, noodle quarters were assigned a label: “A”, “B”, “C”, or “D”. Polymerase Chain Reaction (PCR) amplification of the V4 region of the 16S rRNA gene followed the Earth Microbiome Protocol (https://earthmicrobiome.org/) with the updated 515F (GTGYCAGCMGCCGCGGTAA) (34) and 806R (GGACTACNVGGGTWTCTAAT) (35) primer set. To achieve a dual-indexing sequencing strategy, PCR primers were modified to contain the full P5 and P7 Illumina adapters on the 5’ ends and these adapters contained unique index dual barcode sequences (36). PCR was set up as follows: 2 µL forward and reverse primer mixture (5µM each; 0.4µM reaction concentration), 2 µL support media template DNA, 21 µL PCR-grade molecular water (final volume: 25 µL). This mixture was pipetted into bead tubes containing dried down Cytiva Hot Start Master Mix (Marlborough, MA). Thermocycler conditions were as follows: 95 °C for 3 minutes; 25 cycles of 95 °C for 45 seconds, 50 °C for 45 seconds, 72 °C for 1 minute; 72 °C final extension for 10 minutes. Successful gene amplification was confirmed through gel electrophoresis on a Lonza FlashGel System (Basel, Switzerland). PCR product was purified and pooled by equal volume using the Zymo Research DNA Clean & Concentrator-5 Kit (Irvine, California). Libraries underwent high-throughput sequencing using the Illumina MiSeq platform with 2×250 (V2) paired-end reads at the University of Colorado Anschutz Medical Campus.

### 16S rRNA data analysis

The 16S rRNA data analysis protocols, including specific parameters for each step, can be found at: https://github.com/jessieromero418/Lowry-2025-Paper/tree/main/16S. Raw sequencing data and Phred-33 quality scores were imported into QIIME 2 (Version 2023.5.0) with forward and reverse read files that pertained to each of the 30 available samples from the four sampling timepoints studied. Demultiplexed sequences were denoised within QIIME 2 using the DADA2 denoise-paired plugin (parameters: --p-trim-left-f 0, --p-trim-left-r 0, --p-trunc-len-f 250, --p-trunc-len-r 60, --p-max-ee-f 2, --p-max-ee-r 2, --p-trunc-q 2) and amplicon sequence variants (ASVs) were defined. Taxonomy was assigned to the resulting sequences with the feature-classifier classify-sklearn plugin and the Naïve Bayes classifier trained on the SILVA database (Release 138) (37, 38). To help confirm SILVA taxonomic assignments, the ASV representative sequences underwent a BLASTn search against National Center for Biotechnology Information (NCBI) prokaryote (nt_prok, downloaded May 27, 2024) and virus (nt_viruses, downloaded June 5, 2024) databases. Samples with less than 5300 reads were removed, reducing this dataset to 23 samples across our selected timepoints (2019-03-26, n=5; 2022-01-18, n=7; 2022-01-25, n=6; 2022-03-22, n=5). Additionally, ASVs that were identified as chloroplasts, mitochondria, were unassigned, or were only assigned to the domain level and had no BLAST nr hit, were filtered out of the ASV table. The feature table and representative sequences derived from this filtered dataset were used in downstream analysis. The BIOM table and ASV representative sequences are available in the Supplementary Materials. A taxonomy table was generated from this dataset with the metadata tabulate plugin. A rooted phylogenetic tree was created using the phylogeny align-to-tree-mafft-fasttree plugin.

The resulting feature table, taxonomy table, rooted phylogenetic tree, and metadata file were uploaded to RStudio (Version 4.3.0) with the mia package (Version 1.8.0) (39) for further analyses and figure generation. To shorten taxonomic labels in plots, the BiocParallel (Version 1.34.2) (40), stringr (Version 1.5.0) (41), S4Vectors (Version 0.38.1) (42), stats4 (Version 4.3.0), and BiocGenerics (Version 0.46.0) (43) packages were used to remove rank prefixes (e.g., “p” labels, indicating the phylum level, were removed). The mia scater (Version 1.28.0) (44) and patchwork (Version 1.1.2) (45) packages were used to run and display alpha diversity analyses. The phyloseq package (Version 1.44.0) (46) was used to convert the data, a TreeSummarizedExperiment object, into a phyloseq object. The phyloseq (Version 1.44.0) (46), ggplot2 (Version 3.4.2) (47), and patchwork (Version 1.1.2) (45) packages were used to run and display beta diversity analyses. The miaViz (Version 1.8.0), ggplot2 (Version 3.4.2) (47), and ggraph (Version 2.1.0) (48) packages were used to create taxonomy bar plots of the top 20 phyla and families. At the family level, taxa identified as “uncultured” families and those not in the top 20 families were grouped under the “All other families” category.

### Metagenomic shotgun sequencing

To describe SDIMO genes and visualize gene neighborhoods (spatially co-located genes in a region of a genome), high-throughput, whole-genome shotgun metagenomic sequencing was performed on support media. Support media from each of the four sampling time points (listed above) were processed in triplicates, resulting in a total of 12 samples. Using two quarter sections of a single support media cut with a sterilized razorblade, DNA was extracted with the QIAGEN DNeasy PowerSoil Kit (Hilden, Germany). Bead beating occurred on a laboratory vortex with an adapter for 12 tubes for 10 minutes, and no heating step was added. DNA was eluted in 50 µL of sterile molecular grade water. DNA purity and concentration were assessed using Nanodrop OneC (Waltham, Massachusetts). The University of Colorado Anschutz Medical Campus Genomics Shared Resource (https://medschool.cuanschutz.edu/colorado-cancer-center/research/shared-resources/genomics) prepared libraries using the NuGEN/Tecan Ovation Ultralow System V2 kit and sequenced the DNA using the Illumina NovaSeq 6000 platform with 2×150 paired-end reads on an S4 flow cell lane.

### Metagenomic assembly

The metagenomic shotgun sequencing analysis protocols from assembly to Tier 2 analyses, including specific parameters for each step, can be found at: https://github.com/jessieromero418/Lowry-2025-Paper/tree/main/Shotgun. Raw, forward and reverse read sequencing data files were interleaved with reformat.sh (Version 38.96) (49) for each of the 12 samples. Illumina adapters were trimmed, low-quality bases were trimmed, duplicates were removed, and optical duplicates were removed from interleaved files with rqcfilter2.sh (parameters: rna=f, trimfragadapter=t, qtrim=rl, trimq=0, maxns=3, maq=3, minlen=51, mlf=0.33, phix=t, removehuman=f, removedog=f, removecat=f, removemouse=f, khist=f, removemicrobes=f, clumpify=t, dedupe=t, opticaldupes=t, ddist=12000,tmpdir=, barcodefilter=f, trimpolyg=5, usejni=f) (Version 38.96) (49). A sample of 10,000 resulting trimmed reads were assessed for quality with FastQP (Version 0.3.4) and FastQC (Version 0.11.9) (50). All FastQC results were aggregated to compare read quality across samples with MultiQC (Version 1.12) (51). Trimmed reads were deinterleaved with reformat.sh for downstream analysis. *De novo* assembly of deinterleaved, trimmed reads was performed with MEGAHIT and the --presets meta-large parameter (Version 1.2.9) (52). Each of the 12 samples’ reads were assembled individually. The resulting assemblies were assessed for quality with QUAST (Version 5.2.0) (53). All trimmed read sets were mapped against all assemblies with BBMap and the ambiguous=random parameter (Version 38.96) (49). The resulting 144 BAM files were sorted with samtools sort (Version 1.17) (54). Sorted BAM files required two steps of pre-processing for downstream analysis: 1) contig information was trimmed from sorted BAM files with reformat.sh (Version 38.96) (49) and the parameter trimrname=t, and 2) MD tags were generated in reformatted, sorted BAM files with samtools calmd (Version 1.17) (54). Prodigal (Version 2.6.3) (55) was used with the -p meta parameter to predict gene sequences in the assembly contigs of all 12 samples.

### Homology search for soluble di-iron monooxygenases

To assess the presence of soluble di-iron monooxygenases (SDIMOs) in bioreactor microbial communities, sequences pertaining to the alpha hydroxylase subunit of SDIMOs were searched for in the 12 whole-genome shotgun metagenomic sequencing samples. Contigs of all sizes were analyzed to avoid exclusion of potential plasmid sequences located on smaller contigs. BLASTp (Version 2.9.0) (56) was used to search for SDIMO alpha hydroxylase subunits in proteins predicted by Prodigal. Searches were conducted in protein space to characterize the physico-chemical properties of amino acid residues and describe potential protein functions. A total of 39 recently described SDIMO sequences (15) were used as query sequences against a database of all predicted Lowry proteins at an e-value of 1e-50. These were searched for as they represent a diverse, comprehensive set of candidate dioxane-degrading and non-dioxane-degrading SDIMOs that may aid in the identification of dioxane biodegradation mechanisms. Query sequences belonged to one of four categories: 1) candidate dioxane-degrading proteins (CDDPs) from the literature with evidence of dioxane degradation, 2) outgroup proteins (OUTs) from the literature with no evidence of dioxane degradation, 3) composite proteins (COMPs), which were curated with BLASTp from the genomes of known dioxane-degrading bacteria after searching these for CDDPs, and 4) composite outgroup proteins (COMPOUTs), which were curated similarly to COMPs but were presumed not to degrade dioxane based on phylogenetic placement (15). The 39 SDIMO queries were assigned to Kyoto Encyclopedia of Genes and Genomes (KEGG) Orthology (KO) groups using GhostKOALA (https://www.kegg.jp/ghostkoala/) (57) to predict protein functions. Included among the 39 sequences was a Group IV SDIMO (COMP11, putative alkene monooxygenase alpha subunit [*Mycolicibacterium rhodesiae* JS60] AAO48576.1). Although Group IV SDIMOs have not been associated with dioxane degradation to date (24), this sequence was kept in this analysis for comparison. Also, although one of the 39 sequences lacked both DE*RH motifs of the carboxylate di-iron center (COMP7, soluble di-iron monooxygenase alpha subunit, partial [*Pseudonocardia sp.* D17] BAU36819.1), it was still considered as it was associated with direct metabolism of dioxane (15, 58). Descriptions of the 39 query sequences can be found in Tables S1 and S2.

### Ranking soluble di-iron monooxygenases

A tiered approach was used to rank Lowry protein sequences obtained from BLASTp hits based on levels of evidence of encoding SDIMOs (a flowchart visualizing this approach is shown in Fig. S2). Tier 1 represented protein sequences with the most evidence of encoding SDIMOs, and these were the primary focus of this paper. Tier 1 was curated by filtering BLASTp outputs to only include proteins with a minimum of 90% amino acid identity with a sequence from K. L. Goff and L. A. Hug (15), a minimum alignment length of 125 amino acids, and a minimum query coverage of 60%. This resulted in a total of 86 protein sequences in Tier 1, each originating from a separate contig (Fig. S2). Candidate SDIMOs were further evaluated by aligning the sequences with the 39 SDIMO sequences described by K. L. Goff and L. A. Hug (15) using MAFFT and the –op 1.53 and –ep 0.123 parameters (Version 7.123b) (59). Protein alignments were used to confirm the presence of one or two DE*RH motifs in carboxylate di-iron centers commonly associated with SDIMOs (60). Proteins were included in Tier 1 if they contained at least one DE*RH di-iron center motif.

A phylogenetic tree was generated from the protein alignment using FastTree (61), then annotated in iTOL (62). The tree was re-rooted with Group V SDIMO outgroup sequences. Using monophyletic clades in the tree containing a documented dioxane-degrading sequence, Tier 1 sequences were then categorized as either potential dioxane-degrading proteins or potential outgroup proteins. Clading patterns with previously annotated sequences were also used to aid sequence classification into SDIMO Groups I-VI. To provide more evidence of potential function, proteins were assigned to KO groups using GhostKOALA (57).

Abundances for all 86 Tier 1 sequences were estimated by calculating contig coverages with CoverM (Version 0.6.1) (63) using the reads per kilobase per million mapped reads (RPKM) method. These values were displayed as bar charts in the tree and represent sequence abundances within the sample from which they originated.

To remove identical and near-identical contigs assembled from multiple samples, the contigs from which the 86 Tier 1 protein sequences originated were dereplicated in nucleotide space with CD-HIT-EST (c=0.999, aS=0.999) (Version 4.8.1) (64, 65) (Fig. S2). This resulted in a set of 57 non-redundant contigs, each containing one SDIMO alpha hydroxylase sequence. The SDIMO alpha hydroxylase protein sequences pertaining to the 57 non-redundant contigs were aligned then visualized in a phylogenetic tree with the 39 SDIMO sequences from K. L. Goff and L. A. Hug (15) as described above. To manually cluster the Tier 1 tree and analyze general trends across SDIMOs, the most abundant sequence was identified within each clade (Fig. S2). This sequence was then used as the representative Lowry sequence for each clade. If the most abundant sequence was found to contain a gap in the first di-iron center of the alpha hydroxylase (i.e., the first DE*RH motif was missing in the protein alignment), then the next most abundant sequence that contained this structure was used as the representative Lowry sequence instead. When selecting representatives, the presence of the first di-iron center region was prioritized over the second region as it is near the hydrophobic residues that determine SDIMO group and function (66, 67). Using protein alignments, these hydrophobic residues helped confirm SDIMO classification into Groups I-VI. This reduced the 57 Lowry protein sequences to seven clusters, each with one representative sequence. These seven cluster representative sequences and the 39 SDIMOs from K. L. Goff and L. A. Hug (15) were aligned as described previously and visualized in Microsoft Excel and Jalview (Version 2.11.2.7). A phylogenetic tree was generated from this alignment as described previously. To compare cluster abundances, the abundances of the original 86 Tier 1 protein sequences were summed within each cluster if they originated from the same sample. These summed values were displayed in heatmaps in the clustered tree and represent total abundances of sequences within each cluster within each sample.

To compare sequence diversity, the SDIMO alpha hydroxylases within the 57 non-redundant contigs were clustered in protein space with CD-HIT (Version 4.8.1) (64, 65) and in nucleotide space with CD-HIT-EST (Version 4.8.1) (64, 65) at various sequence identity thresholds (c=1, 0.99, 0.97, 0.95, 0.9).

Tier 2 represented Lowry protein sequences with less evidence of encoding SDIMO alpha hydroxylases. This was curated by filtering BLASTp outputs to include proteins with 50-90% amino acid identity with a sequence from K. L. Goff and L. A. Hug (15), a minimum alignment length of 125 amino acids, and a minimum query coverage of 60%. This resulted in a total of 622 potential Tier 2 SDIMOs that originated from 606 separate contigs (Fig. S2). Contigs that appeared in Tier 1 were removed from this set, resulting in 564 Tier 2 contigs. The 564 Tier 2 contigs were also dereplicated to remove identical and near-identical contigs in nucleotide space with CD-HIT-EST (c=0.999, aS=0.999) (Version 4.8.1) (64, 65) (Fig. S2). This resulted in 355 non-redundant contigs, containing 367 Tier 2 SDIMOs. The 367 Tier 2 protein sequences and the 39 SDIMO sequences described by K. L. Goff and L. A. Hug (15) were aligned with MAFFT and the –op 1.53 and –ep 0.123 parameters (Version 7.123b) (59). An unrooted phylogenetic tree was generated from the protein alignment using FastTree (61), then annotated in iTOL (62). Tier 2 sequences were colored by predicted SDIMO group based on clading patterns with literature sequences.

### Gene neighborhoods surrounding soluble di-iron monooxygenases

The gene neighborhoods associated with the Lowry contigs containing Tier 1 Group V, DxmA SDIMO alpha subunit genes (n=17 contigs) were analyzed. All 17 contigs were annotated in the DOE Systems Biology Knowledgebase (KBase) (68) with the “Annotate and Distill Assemblies with DRAM” (v0.1.2) application (69) and the “Annotate Metagenome Assembly and Re-annotate Metagenome with RASTtk” (v1.9.5) application (70). This provided gene annotations from DRAM (71), RAST (72), KEGG (73), and the Protein Families Database (PFAM) (74). Predicted proteins were also analyzed using a protein BLAST against the NCBI non-redundant database (56). For protein BLAST results, the annotation with the lowest E-value was selected, except in cases where the protein was classified as hypothetical or unknown. In these instances, the next lowest E-value hit with an annotated protein was considered, provided the E-value remained below 1e^-05^. Final annotations were assigned when two or more of the annotation methods (DRAM, RAST, KEGG, PFAM, BLASTp against nr) agreed on the same annotation. In instances where none of the annotations agreed, the BLASTp result was taken (as long as the E-value < 1e^-05^) and the annotation was named as ‘putative’. If no annotations were identified in any of the methods, the protein was called ‘hypothetical’. To gather more evidence of potential horizontal gene transfer, the 17 contigs were evaluated for plasmid and viral sequences using geNomad (Version 1.7.0) with default parameters (75). Putative transposase gene sequences were evaluated with TnCentral using default parameters (76). Putative toxin/antitoxin gene sequences were evaluated with TAfinder 2.0 using default parameters (77). Putative miniature inverted-repeat transposable elements (MITEs) were evaluated with MITE Tracker using default parameters (78).

The 17 DxmA contigs were aligned using Geneious Prime (Version 2024.0.7; https://www.geneious.com) with the ‘map to reference’ feature utilizing the Geneious mapper on the highest sensitivity using “find structural variants, short insertions, and deletions of any size”, with contig S3_k127_1715793 as the reference sequence. Four contigs containing candidate dioxane-degrading proteins identified in K. L. Goff and L. A. Hug (15) were also aligned with the Lowry contigs. Phylogenetic analysis was done using RAxML Version 4 (79) with a GTR GAMMA nucleotide model and a rapid hill-climbing algorithm and 100 bootstrap replicates. One additional reference sequence (*Pseudonocardia asaccharolytica* DSM 44247; NZ_AUII01000002) trimmed to only include the dioxane monooxygenase alpha, beta, coupling, and reductase components) was
used to root the tree.

### Data availability

Reads have been deposited to the NCBI SRA database under BioProject PRJNA1233214.

## RESULTS

### Site chemistry

To characterize the microbial community and genomically encoded metabolic potential of the BTS, genomic DNA was extracted from biofilms growing on independent support media samples taken on each of four sampling dates. Site operators regularly take measurements of key chemical concentrations flowing into the BTS. For the days sampled, input dioxane concentrations ranged from 15,000 µg/L to 28,000 µg/L, while the median of all weekly measurements taken in the 3-year sampling period was 19,000 µg/L (Fig. 1). On average, BTS effluent dioxane levels measure less than 25 µg/L. THF was also monitored since some SDIMO enzymes are known to co-metabolize dioxane alongside THF. THF concentrations in the BTS influent ranged from 16,000 µg/L to 31,000 µg/L on the days sampled (median in time period = 21,000 µg/L).

**Fig 1.**
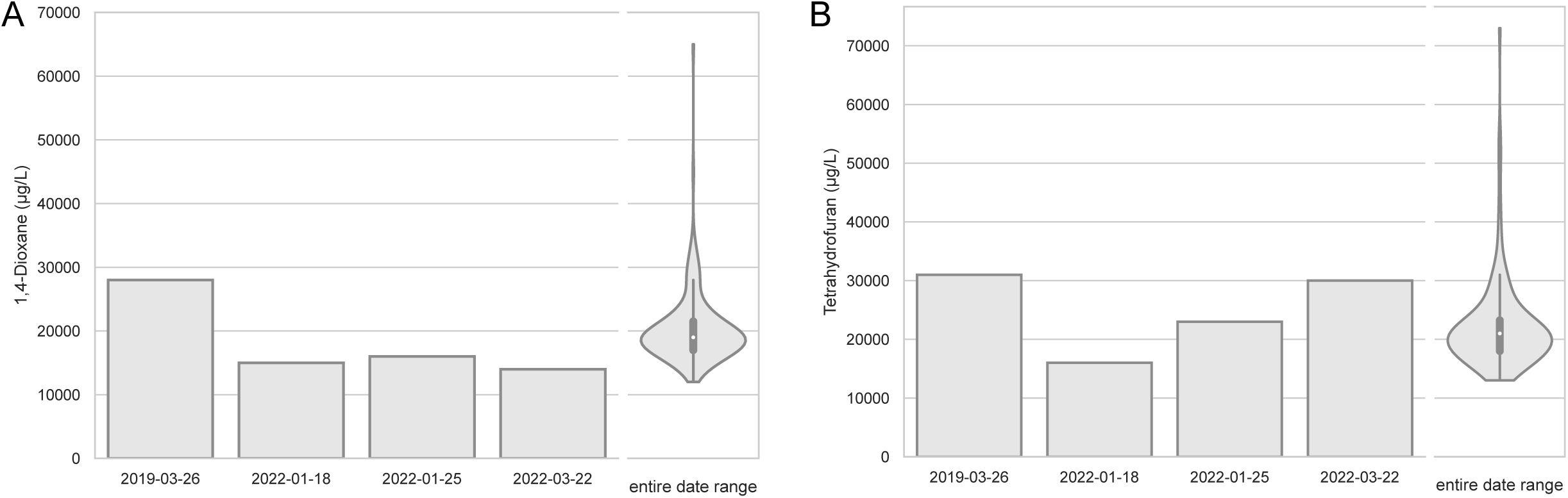
Input dioxane concentrations to the sedimentation tank of the Lowry Landfill Biological Treatment System (BTS). Values for each of the dates sampled for shotgun metagenomics are shown alongside the distribution (kernel density estimate and quartiles / whiskers) of values from regular weekly sampling between the first and last sampling date. The median concentration for dioxane weekly measurements over the three-year sampling date range was 19,000 µg/L (A). The median concentration for THF, a common substrate for SDIMO enzymes that co-metabolize 1,4-Dioxane, was 21,000 µg/L (B).

### Microbial community composition of bioreactor support media

To describe the microbial community composition of Bioreactor 1, 16S rRNA gene sequences were analyzed from 23 Bioreactor 1 support media samples (Table S3) over the selected timepoints (2019-03-26, n=5; 2022-01-18, n=7; 2022-01-25, n=6; 2022-03-22, n=5) A total of 884 ASVs were identified across this dataset. Alpha diversity analyses across the samples showed Observed Richness ranging from 113-508, evenness (Shannon metric) from 3.8-4.5, Simpson metric from 0.04-0.20, and phylogenetic diversity (Faith PD metric) from 11-36 (Fig. S3, Table S3). Non-metric multidimensional scaling (NMDS) beta diversity analyses based on Unifrac distances showed overall community stability across samples and time points (Fig. S4). The 2019 samples had somewhat different richness and diversity estimates from the 2022 samples and clustered independently in the NMDS plots but overall had similar taxonomic groups. This suggests that the overall community membership was similar, but there were some slight differences in community structure. Overall, the microbial communities were very stable over time.

Microbial community composition was diverse across multiple taxonomic levels. A total of 31 phyla were identified across all samples. At the phylum level, communities were consistently dominated by Proteobacteria (34.1-53.5%), followed by Nitrospirota (7.4-23.9%), Actinobacteriota (5.5-17.5%), and Crenarchaeota (4.7-15.6%) (Fig. 2A). Other commonly occurring phyla included Acidobacteriota, Planctomycetota, Bacteroidota, Verrucomicrobiota, Myxococcota, and Chloroflexi (Fig. 2A). At the family level, approximately 196 families were identified in the bioreactor samples. The most abundant families included Nitrospiraceae (7.3-23.8%), Nitrososphaeraceae (4.6-15.6%), Nitrosomonadaceae (4.8-13.2%), Pseudonocardiaceae (1.8-8.4%), Hyphomicrobiaceae (2.6-6.3%), Parvularculaceae (1.9-7.5%), Steroidobacteraceae (3.2-5.4%), Saprospiraceae (1.7-3.8%), Rhodocyclaceae (1.1-4.3%), and Methyloligellaceae (0.4-3.4%) (Fig. 2B). Overall, the taxonomy showed stability over time and across individual samples.

**Fig 2.**
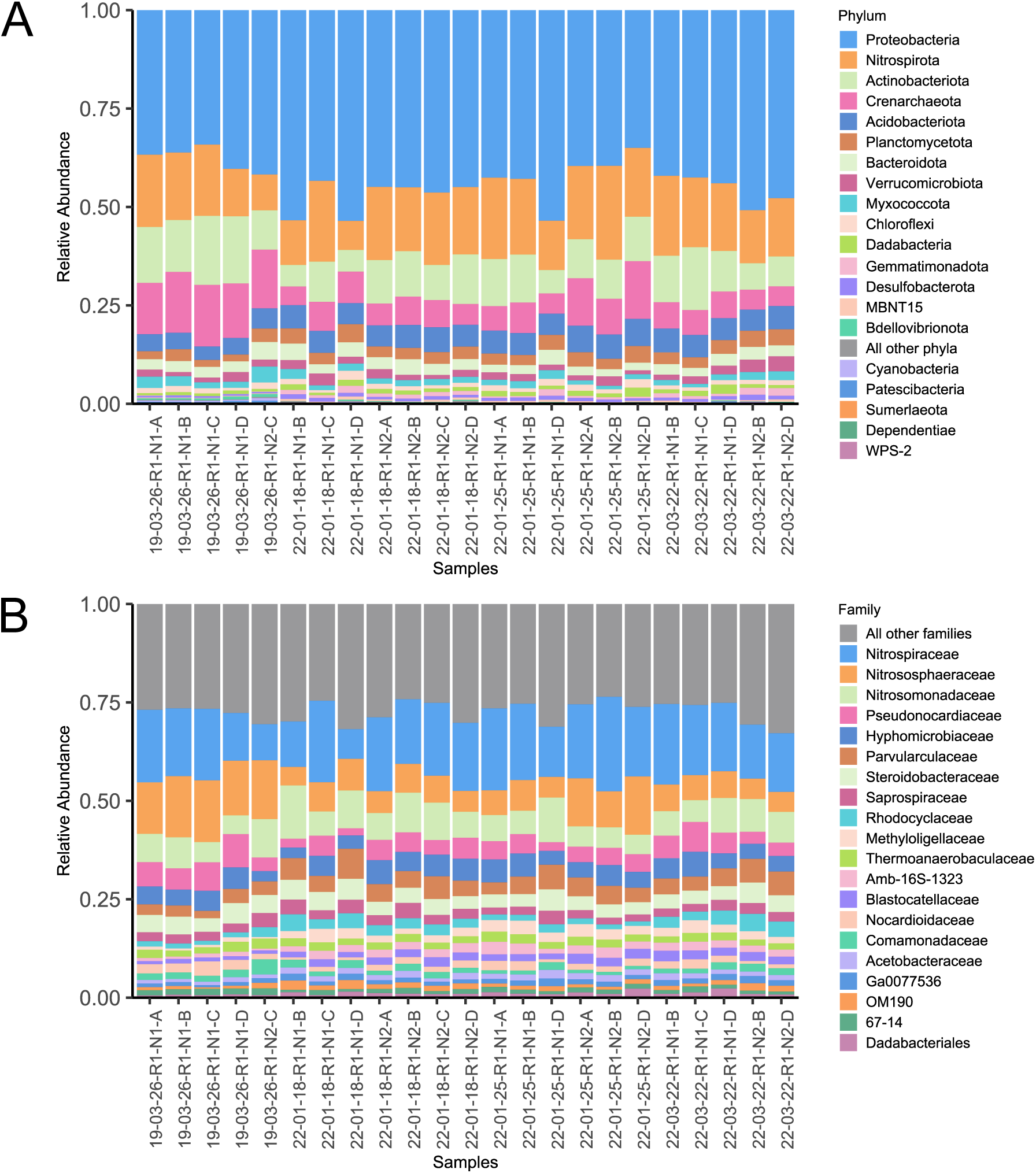
Microbial community composition bar plot of the top 20 most abundant phyla (A) and families (B) across Lowry Landfill Bioreactor 1 support media (n=23). “All other phyla” represent phyla that were not among the top 20 most abundant phyla. “All other families” represent families that were not among the top 20 most abundant families and families that were only identified as “uncultured”.

Many of the ASVs in the dataset represent potential novel groups: 150 were identified as “uncultured” at the genus level (with no genus identified), 71 were identified only down to the family level (with no genus), 10 identified only to the order level, 18 identified only to the class level, 9 identified only to the phylum level, and 29 identified only to the domain level.

Several ASVs were taxonomically related to bacterial genera containing previously reported dioxane degraders: 8/884 ASVs were identified as *Pseudonocardia* (two of these ASVs were present in 23/23 samples), 4/884 as *Mycobacterium* (one of these ASVs was present in 23/23 samples), 2/884 as *Flavobacterium* (present in 4/23 samples total), 2/884 as *Acinetobacter* (present in 2/23 samples total), 3/884 as *Pseudomonas* (present in 6/23 samples total), and 1/884 as *Afipia* (present in 5/23 samples total). These results suggest that bacteria that were closely related to known dioxane degraders were present across Lowry support media.

### SDIMO diversity and abundance

To describe the bacterial proteins potentially involved in dioxane degradation in Bioreactor 1, predicted proteins on Lowry contigs assembled from 12 shotgun sequencing samples (Table S4) were searched against 39 known SDIMO alpha hydroxylases (15) using BLASTp. This revealed a phylogenetically diverse set of 86 Tier 1 SDIMO alpha hydroxylases in the support media. After clustering the contigs containing the 86 SDIMO alpha hydroxylases, a non-redundant set of 57 contigs was obtained (Fig. S2). SDIMOs pertaining to Groups I (n=2), II (n=4), IV (n=7), and V (n=73) were observed (Fig. S5). Three SDIMO groups of candidate dioxane-degrading proteins (CDDPs) were identified based on clading patterns, each of which was assigned one representative Lowry sequence (Fig. 3): Group II CDDPs (n=3), Group V, PrmA*-like CDDPs (n=13), and Group V, DxmA-like CDDPs (n=38). Additional protein sequences phylogenetically grouped with a Group IV SDIMO (a putative alkene monooxygenase, n=7), as well as outgroups for Group I (n=2), Group II (n=1), and Group V, PrmA*-like outgroups (n=22). Group I sequences were assigned to K16242 (dmpN, poxD, tomA3; phenol/toluene 2-monooxygenase (NADH) P3/A3) and Group II sequences to K15760 (tmoA, tbuA1, touA; toluene monooxygenase system protein A). In Group IV, 6/7 sequences were assigned to K18223 (prmA; propane 2-monooxygenase large subunit) with a second possible assignment to K22353 (etnC, alkene monooxygenase alpha subunit). One Group IV sequence (LOWRY-S8_k127_2122582_2) lacked a KO assignment. Group V sequences were assigned to K18223. Although previous studies have found CDDPs pertaining to Group III (K16157: mmoX; methane monooxygenase component A alpha chain) (18) and Group VI (80), no Lowry proteins curated in this protocol were identified as such. Thus, Tier 1 included all SDIMO groups, except Groups III and VI.

**Fig 3.**
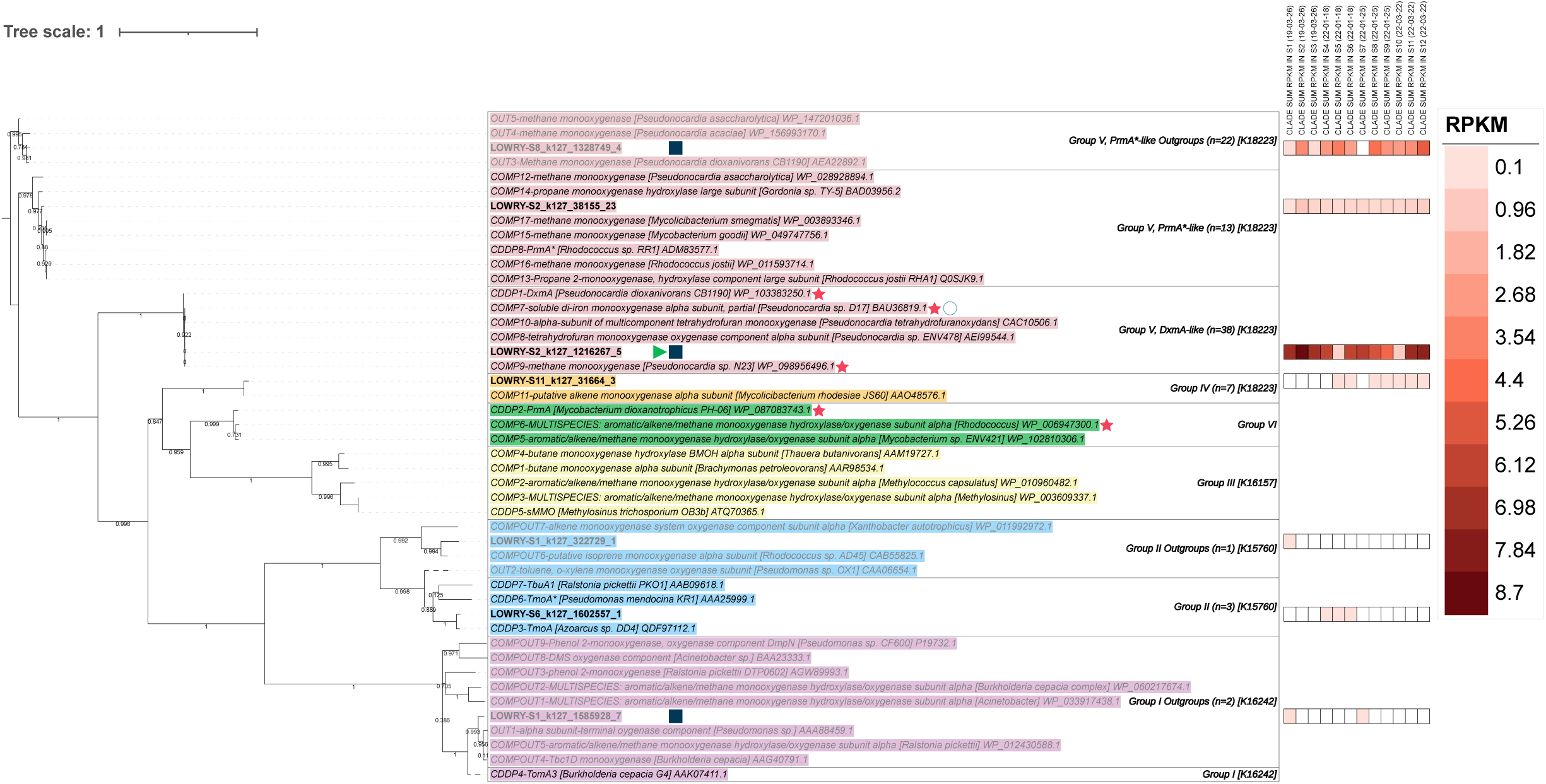
Manually clustered protein phylogenetic tree of seven representative sequences recovered from Lowry Landfill Bioreactor 1 support media and 39 candidate SDIMOs described by K. L. Goff and L. A. Hug (15). Protein abundances were estimated using coverages (RPKM) of the 86 original Lowry contigs containing SDIMOs, summed by clade and by sample, and are displayed in the heatmap. Proteins that are presumed not to degrade dioxane due to monophyletic clading with known outgroups are written in gray text. The branches for OUT2 and CDDP4 were dashed as these showed monophyletic clading that was unexpected for their description in K. L. Goff and L. A. Hug (15). Red stars indicate sequences that have shown direct metabolism of dioxane in the literature. Empty circles indicate sequences containing a gap in the first di-iron center of the SDIMO alpha hydroxylase (i.e., missing the first DE*RH motif). Clades that contained at least one sequence with an inverted terminal repeat or a plasmid score above the default threshold according to geNomad were marked with green triangles or blue squares, respectively. The SDIMO group number, number of sequences per phylogenetic clade, and KO assignments of representative sequences are listed in clade boxes.

To compare the prevalence of these proteins, abundances were estimated using contig coverage (RPKM) associated with the 86 Tier 1 sequences (Fig. 3). Within each monophyletic clade, the most abundant contig that did not show a gap in the first di-iron center of the alpha hydroxylase was chosen as a representative and displayed in the tree. SDIMO classification was confirmed for these sequences in protein alignments using hydrophobic residues surrounding the first di-iron center region (see Methods and Fig. S6). When abundances were summed across all 12 samples by group, Group V sequences showed the highest total abundance (105.4 RPKM), followed by Group IV (1.1 RPKM), Group II (0.5 RPKM), and finally Group I (0.3 RPKM). Group V sequences were present across all 12 samples. DxmA-like CDDPs were also found in all 12 samples and were the most abundant clade. DxmA-like summed abundances ranged from 0.7-8.7 RPKM per sample and had a total abundance of 68.1 RPKM. The single most abundant Lowry sequence was LOWRY-S2_k127_1216267_5 (6.4 RPKM). This was closely related to DxmA-like CDDPs and was selected as the representative sequence for this clade (Fig. 3). This shared 99% amino acid identity with DxmA from known dioxane-degrading strain CB1190 (WP_103383250.1, CDDP1), and 100% amino acid identity with a partial SDIMO alpha hydroxylase subunit in *Pseudonocardia* sp. D17 (BAU36819.1, COMP7) and a methane monooxygenase from *Pseudonocardia* sp. N23 ([WP_098956496.1](https://www.ncbi.nlm.nih.gov/protein/BAU36819.1), COMP9). All three of these proteins have shown direct metabolism of dioxane (15, 18, 58, 81, 82). The LOWRY-S2_k127_1216267_5 sequence also shared high amino acid identity with DxmA-like sequences that showed co-metabolic dioxane degradation in the presence of THF (15, 83). For example, this shared 99.8% identity with the alpha hydroxylase in *Pseudonocardia* sp. ENV478 (AEI99544.1, COMP8). Overall, Group V was the most abundant SDIMO group in Tier 1, with certain proteins sharing high amino acid identity to directly metabolic and co-metabolic DxmA-like CDDPs.

Because several Lowry sequences claded with DxmA-like CDDPs, these were analyzed further to describe the protein and gene sequence diversity of this clade. Clustering the alpha hydroxylases within the 57 non-redundant contigs resulted in multiple DxmA-like CDDP representatives, even at various stringency levels of sequence similarity (Table 1). For example, after clustering the 57 alpha hydroxylase proteins from the non-redundant contigs at 100% amino acid identity, 38 non-redundant proteins were obtained. Of these proteins, 14 claded with DxmA-like CDDPs. After clustering the 57 proteins at 90% amino acid identity, 2/10 remaining proteins claded with DxmA-like CDDPs. At the nucleotide level, clustering the 57 alpha hydroxylase genes at 100% nucleotide identity resulted in 45 non-redundant genes. Of these, 15 were identified as DxmA-like genes. Clustering at 90% nucleotide identity resulted in 19 genes, 7 of which were identified as DxmA-like genes (Table 1). DxmA-like protein and gene sequences were consistently clustered into multiple groups, illustrating the amino acid and nucleotide sequence diversity of this one clade.

**Table 1.** SDIMO alpha hydroxylase subunit gene sequence clustering within 57 non-redundant Tier 1 contigs.

| Clustering Parameters |  | Protein-level Clustering |  | Nucleotide-level Clustering |  |
| --- | --- | --- | --- | --- | --- |
| c | aS | Total SDIMOs | Number of DxmA sequences | Total SDIMOs | Number of DxmA sequences |
| 1 | NA | 38 | 14 | 45 | 15 |
| 0.99 | NA | 30 | 9 | 32 | 11 |
| 0.97 | NA | 24 | 7 | 28 | 10 |
| 0.95 | NA | 22 | 7 | 24 | 9 |
| 0.9 | NA | 10 | 2 | 19 | 7 |

CD-HIT clustering parameters: c = percent identity, aS = alignment coverage for the shorter sequence (64, 65).

Other CDDPs identified in the Lowry samples included Group V, PrmA*-like CDDPs, represented by LOWRY-S2_k127_38155_23 (Fig. 3). This shared 96.4% amino acid identity with CDDP8, the co-metabolic PrmA* protein identified in *Rhodococcus* sp. RR1 (ADM83577.1) that was associated with dioxane degradation after induction with toluene (18). Other co-metabolic SDIMOs found in this clade included PrmA*-like proteins from *Rhodococcus jostii* RHA1 (Q0SJK9.1, COMP13; WP_011593714.1, COMP16), a bacterium that degraded dioxane in the presence of propane and 1-butanol growth substrates (84). PrmA*-like CDDPs were detected in all 12 samples at relatively low summed abundances (0.1-1.1 RPKM per sample) and a total abundance of 6.4 RPKM. Group II CDDPs were also detected. These were represented by LOWRY-S6_k127_1602557_1 (Fig. 3), which claded and shared 95.8% amino acid identity with CDDP3, the co-metabolic TmoA protein identified in *Azoarcus* sp. DD4 (QDF97112.1) that degraded dioxane after induction with toluene (85). Group II CDDPs were only detected in a subset of the samples at low summed abundances (0.1-0.2 RPKM per sample, total abundance=0.4 RPKM). Lastly, Group IV SDIMOs were detected. However, these have not been associated with dioxane degradation to date (24). These sequences were represented by LOWRY-S11_k127_31664_3 (Fig. 3), which shared 96.6% amino acid identity with a putative alkene monooxygenase alpha subunit in *Mycolicibacterium rhodesiae* JS60 (AAO48576.1, COMP11). Group IV SDIMOs were detected in a subset of the samples at low summed abundances (0.1-0.2 RPKM per sample, total abundance=1.1 RPKM). The presence of these SDIMOs provides evidence for other potential dioxane co-metabolism mechanisms, and for biodegradation of other chemicals in the Lowry bioreactor.

### Initial evaluation of potentially novel SDIMOs

Using less strict criteria, Tier 2 was developed from Lowry Landfill Bioreactor 1 proteins and represented those with less evidence from sequence homology of encoding SDIMO alpha hydroxylases. Tier 2 consisted of 622 potential SDIMOs. After filtering out Tier 1 contigs, clustering resulted in 355 non-redundant contigs containing a total of 367 potential SDIMOs (Fig. S2). An unrooted protein phylogenetic tree of the 367 potential SDIMOs and the 39 SDIMOs from K. L. Goff and L. A. Hug (15) revealed a phylogenetically diverse set of sequences predicted to pertain to Groups I (n=130), II (n=169), III (n=1), IV (n=18), V (n=42), and VI (n=7) (Fig. S7). The Tier 2 potential SDIMO proteins were closely related to diverse CDDPs: Group I TomA3-like CDDPs, Group II CDDPs, Group III sMMO-like CDDPs, Group V PrmA*-like CDDPs, and Group VI PrmA-like CDDPs. Therefore, this protocol curated additional, more phylogenetically diverse SDIMOs that were in the bioreactor. Further efforts are needed to determine the function of these proteins, but they may represent novel, uncharacterized SDIMOs involved in dioxane or other co-contaminant biodegradation.

### Search for evidence of dioxane degradation pathway steps and horizontal gene transfer

We sequenced and assembled Lowry contigs to evaluate the SDIMOs and CDDPs in a genomic context. However, in closely evaluating the sequence data, we noticed that the assembly was breaking across several of the relevant contigs. For instance, assembly breaks were noted in the S3_k127_1715793 and S2_k127_1904549 contigs, which were each longer than 27,000 nucleotides and contained abundant Group V, DxmA-like proteins (Fig. 4). An additional 20 shorter contigs mapped to these longer contigs with high identity (~99% nucleotide identity across all contigs), including contigs from all samples except for S9. Curiously, there were four separate regions of overlap in the alignment where the assembly appeared to break. The four sequence overlaps were each exactly 127 bp long, but each contained different nucleotide sequences, suggesting micro-variation in the population. Three out of the four sequence overlaps occurred near the end of a protein-coding gene.

**Fig 4.**
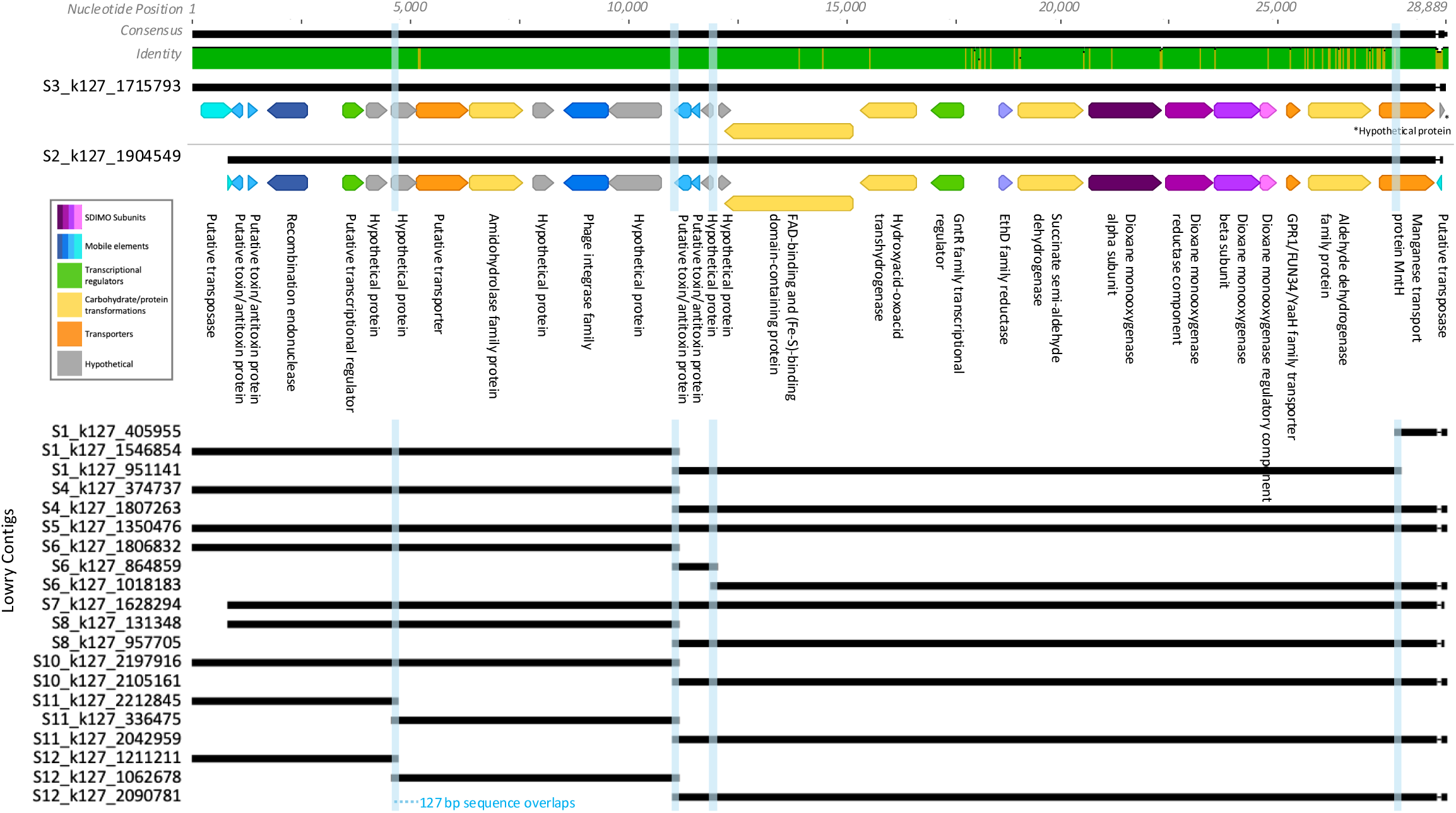
Nucleotide alignment of overlapping contigs. Blue columns indicate the 127 base pair stretches where the assembly broke. Predicted proteins and annotations (color coded by function; see inset) are shown for the two longest contigs.

Given the challenges with genome assembly, our analysis of SDIMO gene neighborhoods focused on contigs (instead of metagenome-assembled genomes). Based on their high abundance in the Lowry system, we further evaluated the DxmA-like CDDP-containing contigs (n=17 out of the 57 non-redundant contigs) to look for phylogenetic relationships to other known dioxane degraders, evidence of other SDIMO components, dioxane degradation pathway steps, and horizontal gene transfer (Fig. 4, Fig. S8).

Candidate dioxane-degrading contigs identified in K. L. Goff and L. A. Hug (15) were compared to the Lowry contigs. The Lowry contigs were phylogenetically related to the *Pseudonocardia* contigs with strong bootstrap support (>60 for most of the contigs) including: *Pseudonocardia* sp. N23 (NZ_BEGX01000008), *Pseudonocardia* sp. ENV478 (HQ699618), *Pseudonocardia tetrahydrofuranoxydans* strain K1 (AJ296087), and *Pseudonocardia dioxanivorans* EUR-1 (NZ_SJXF01000101). One additional sequence was used as an outgroup by extracting just the SDIMO subunits: *Pseudonocardia asaccharolytica* DSM 44247 (NZ_AUII01000002). This contig-level phylogeny aligned with the protein-level phylogeny showing the individual Lowry DxmA-like alpha subunit proteins phylogenetically clustering with *Pseudonocardia* alpha subunit proteins (Fig. S5).

The gene neighborhood analysis showed that the SDIMO subunit gene order was conserved across nearly all 17 of the Lowry DxmA-like CDDP-containing contigs, only excluding contigs that did not extend over the entirety of the subunits. For each of the complete contigs (containing each of the subunits), the SDIMO subunits were ordered as dioxane monooxygenase alpha subunit, reductase component, beta subunit, and regulatory/coupling component, which has been previously observed in Group V SDIMOs (85). Outside of the SDIMO subunits, predicted semi-aldehyde dehydrogenase, EthD domain, and GntR family transcriptional regulator proteins were located upstream. EthD is a component of a cytochrome P-450 gene, which was associated with the degradation of ethyl tert-butyl (ETBE) in *Rhodococcus rubber* IFP 2001 (86). Downstream of the SDIMO subunits were predicted GPR1/FUN34/yaaH family transporters. Putative proteins occurred at the ends of the contigs, including putative metal transporters and putative transposases. Contigs from K. L. Goff and L. A. Hug (15) had the exact same order of SDIMO subunits as Lowry contigs, and nearly the same upstream and downstream proteins. The contig from *Pseudonocardia dioxanivorans* (NZ_SJXF01000101) extended further downstream than any of the Lowry contigs including a Nramp family divalent metal transporter in place of the putative metal transporters seen in the Lowry contigs, followed by an IS1634 family transposase, tyrosine-type recombinase/integrase, zeta toxin family protein, and two hypothetical proteins.

The Lowry DxmA-like contigs had several genes commonly associated with horizontal gene transfer. Several of these contigs (13/17) were assigned a plasmid identification score above the default threshold (0.7) by geNomad (75) (Fig. S5). Toxin-antitoxin proteins were predicted in the annotated Lowry contigs (S3_k127_1715793, S2_k127_1904549, and S9_k127_1368382). Specifically, TAfinder 2.0 (77) identified a set of Type II toxin-antitoxin phd-doc (prevent host death/death on curing) family proteins, which have been proposed to be involved in plasmid stabilization and many other functions such as stress modulation and antibiotic persistence (87, 88). Other annotation methods (PFAM, BLASTp) identified an additional set of Type II toxin-antitoxin proteins (though not identified by TAfinder 2.0).

According to geNomad, no Lowry proteins had a virus score above the default threshold (0.7), though annotation methods did identify phage integrase proteins likely related to tyrosine family integrases that utilize a catalytic tyrosine to mediate strand cleavage (89).

Putative transposases in the Lowry contigs (S3_k127_1715793_29, S9_k127_1368382_1, and possibly the truncated S2_k127_1904549_1) appeared to be related to the IS5 family of insertion sequences based on BLASTp results (against nr) and TnCentral (76), which have been shown to cause mutations in *E. coli* and are found in taxonomically diverse microbes (76). The S2_k127_1904549 contig had predicted transposases at beginning and end of the contig (S2_k127_1904549_1 and S2_k127_1904549_29), which were 100% identical but in reverse complement orientation (geNomad identified this region as an Inverted Terminal Repeat; Fig. S5). The S3_k127_1715793 contig was dissimilar and had a hypothetical protein at the end of the contig instead of another transposase. Complete insertion sequences or transposons were not identified in any of the Lowry contigs.

Putative miniature inverted-repeat transposable elements (MITEs; short, non-autonomous transposable elements that can contribute to genetic variability and regulation) sequences were identified in the longest Lowry DxmA-like CDDP contigs (S3_k127_1715793, S9_k127_1368382, and S2_k127_1904549) using MITE Tracker (78). The three contigs each had many predicted MITEs across the length of the sequence. Many of the MITEs were overlapping, so the total number of predicted MITEs was uncertain. MITEs were predicted within protein-coding sequences and within intergenic regions. Two of the four 127 bp overlapping regions within the broken assembly (Fig. 4) were within predicted MITEs. The *Pseudonocardia* contigs (NZ_SJXF01000101.1, HQ699618.1, AJ296087.1, NZ_BEGX01000008.1) also contained several MITEs, though fewer than the Lowry contigs in the regions that aligned. The *Pseudonocardia* and Lowry contigs shared a few predicted MITEs with similar structure and genomic location. Further research is needed to confirm whether or not the predicted MITEs are functional.

## DISCUSSION

The microbial communities of the Lowry Landfill Bioreactor 1 are diverse and stable over the sampled three-year period and selected shorter timescales. Our 16S rRNA gene sequence data shows that a number of taxa fall within genera that also contain known dioxane degraders (e.g., *Pseudonocardia*). Also, Pseudonocardiaceae and Rhodocyclaceae, families detected in our system, each contain species that have shown either direct metabolism or co-metabolism of dioxane (15, 16). Taxa in the Hyphomicrobiaceae and Nocardiodaceae families are present in the bioreactor and were previously shown to increase in abundance during dioxane degradation in a microcosm experiment (27). Although the potential Lowry dioxane-degraders were detected in support media, they were not the dominant taxa. Instead, other taxa within the Nitrospiraceae, Nitrososphaeraceae, and Nitrosomonadaceae groups were highly abundant in the support media communities, suggesting an important role for nitrogen cycling. Overall, the 16S data provide first insights into a complex but relatively stable community within the bioreactors. The relationship between dioxane degradation and the other important metabolisms (e.g., nutrient cycling, degradation of other known contaminants in the reactor) in the community remains to be discovered.

From the Tier 1 SDIMO protein dataset, Group V SDIMOs dominated in the Lowry Landfill bioreactor support media bacterial communities. Protein phylogenetic trees and alignments showed Group V SDIMOs consisting of two CDDPs: DxmA-like proteins and PrmA*-like proteins. Of these, the DxmA-like proteins were present across all Lowry support media microbiomes usually as the most abundant SDIMO protein. Some of these proteins shared high amino acid sequence identity with known DxmA-like proteins, including those from ***Pseudonocardia dioxanivorans* str. CB1190** (WP_103383250.1, CDDP1), *Pseudonocardia* sp. D17 (BAU36819.1, COMP7), and *Pseudonocardia* sp. N23 (WP_098956496.1, COMP9), all of which have shown direct metabolism of dioxane (15, 18, 58, 81, 82). Phylogenetic trees showed that some co-metabolic enzymes requiring induction by THF to degrade dioxane, for example, an SDIMO from *Pseudonocardia* sp. ENV478 (AEI99544.1, COMP8) (15, 83), claded with DxmA-like directly metabolic enzymes and Lowry proteins (Fig. 3). Therefore, it is possible that some Lowry proteins in this clade similarly require THF to degrade dioxane co-metabolically. In the BTS, THF concentrations are being monitored as it has been a suspected growth substrate for co-metabolic dioxane degradation. However, the presence of THF-induced co-metabolic degradation of dioxane in the BTS or in the dioxane plume surrounding the facility has not been tested to date. Furthermore, studies have not been able to definitively identify gene sequences or components resulting in either direct metabolism or co-metabolism of dioxane (15). Therefore, Lowry DxmA-like sequences could not be searched for signatures of THF co-metabolism. As a result, although DxmA-like proteins were frequently detected in support media, further experiments are necessary to test not only their dioxane degradation ability, but also their potential degradation mechanism.

The DxmA-like contigs contained many genomic signatures of potential mechanisms for horizontal gene transfer (HGT), including toxin-antitoxin systems, phage integrases, recombination endonucleases, putative transposases, and putative miniature inverted-repeat transposable elements (MITEs). The large number of HGT signatures and diverse mechanisms of function (e.g., plasmids, transposition) within genomic proximity to the SDIMO protein coding genes suggests that these genes may be mobile within the community. The functional consequences of these potential HGT signatures are unknown but could play a role in the microbial evolution within the reactors, population-level variability, and regulation of these important functional genes. For instance, MITEs have been shown to play a role in genome plasticity and gene regulation in bacteria (90, 91). The environmental context of the Lowry bioreactors (a complex contaminant mixture) may put a selection pressure on the community to move the SDIMO genes within the population in order to maintain functional stability. These findings would be interesting to follow up on in the context of dioxane remediation and regulation.

Group V, PrmA*-like proteins were also frequently detected in the Lowry Landfill bioreactor support media, although at lower abundances than the DxmA-like proteins. These shared high amino acid sequence identity with PrmA*, which was associated with co-metabolic degradation induced by toluene in *Rhodococcus sp.* RR1 (18). Other co-metabolic PrmA*-like proteins in this clade were induced by propane and 1-butanol (Q0SJK9.1, COMP13; WP_011593714.1, COMP16) (84). Although propane and 1-butanol are not specifically monitored in the BTS, toluene is measured in BTS effluent. However, whether any of these chemicals induce co-metabolic dioxane degradation by PrmA*-like enzymes in the bioreactors remains an open question. Because these proteins were detected to a lesser extent, it is possible that dioxane degradation by PrmA*-like enzymes is a less common mechanism in the support media bacterial communities.

Compared to Group V, other SDIMO groups were detected more infrequently and to a much lesser degree. Of these, Group II SDIMOs were detected at three orders of magnitude below the Group V SDIMOs. Some Group II SDIMOs claded with the CDDP TmoA, which was induced to degrade dioxane co-metabolically with toluene in *Azoarcus sp.* DD4 (QDF97112.1; CDDP3) (85). Whether dioxane co-metabolism induced by toluene and mediated by TmoA-like enzymes is present in the BTS has not been tested. Additionally, Group IV SDIMOs were detected, although only at a total abundance that was two orders of magnitude lower than that of the Group V SDIMOs. Although Group IV SDIMOs have not been shown to degrade dioxane to date, these proteins have been detected in dioxane-impacted groundwater surrounding industrial sites in Arizona (24) and Alaska (25).

Interestingly, Tier 1 Lowry proteins, those that we were most confident encoded SDIMOs, contained neither Group III nor Group VI SDIMOs. A study on another dioxane-impacted landfill detected sMMO, a Group III SDIMO associated with methane-induced co-metabolic dioxane degradation in OB3b (18), in most landfill monitoring wells by qPCR (92). In the Lowry Landfill, it is possible that bioreactor conditions caused a lack of sMMO-like proteins. For example, sMMO is often detected under copper-limited conditions, and may be replaced by particulate MMO (pMMO) in methanotrophs when copper is not limited (93). The lack of Group VI SDIMOs, like PrmA, is also of note. Prm is a directly metabolic dioxane-degrading enzyme first identified in *Mycobacterium dioxanotrophicus* PH-06 (WP_087083743.1) (80). Compared to the Dxm of CB1190, Prm has shown a higher dioxane degradation rate, broader substrate range, and less susceptibility to inhibition (94).

Tier 2 SDIMOs showed broader phylogenetic diversity than Tier 1. Tier 2 uncovered potential SDIMOs that were closely related to CDDPs not seen in Tier 1, namely Group I TomA3-like CDDPs, Group III sMMO-like CDDPs, and Group VI PrmA-like CDDPs. This illustrates other possible CDDPs in the Lowry Landfill system that could later be studied for dioxane degradation ability.

Several studies have attempted to correlate the presence of different SDIMOs with dioxane degradation ability of bacterial communities. The current study found a potential dominance of DxmA-like proteins in the Lowry Landfill bioreactor support media. This contrasted slightly with the findings of K. L. Goff and L. A. Hug (15), which predicted that DxmA was rare and largely restricted to terrestrial environments while engineered environments were expected to show higher abundances of Tmo (Group II), Tom (Group I), and Tbu (Group II). However, in dioxane-impacted groundwater in Alaska, DxmA-like proteins were detected throughout a dioxane plume but were most prevalent near the source of contamination. Certain Group I SDIMOs were also detected at the source (25). Another study enriched dioxane-degrading microbial communities from two uncontaminated soils. Both soils began with diverse communities and SDIMOs, which were then dominated by *Mycobacterium* and Group V and VI SDIMOs after dioxane addition (95). Also, a study attempting to enrich dioxane-degrading consortia found that Group V, PrmA*-like and Group I, TomA3-like proteins were most abundant in both uncontaminated and contaminated soils. Although there was no statistically significant difference in the abundances of these proteins before and after dioxane addition, *Mycobacterium* was among one of the taxa enriched after dioxane addition (27). Like the current study, another used shotgun sequencing to characterize the SDIMOs and microbial communities of chlorinated solvent-impacted sites. However, these sites were bioaugmented with SDC-9 (Shaw Environmental Inc., Lawrenceville, New Jersey), a dechlorinating bacterial consortium, and one site was confirmed to show dioxane contamination. Across those systems, the most abundant SDIMOs belonged to Groups III, I, then II. *Pseudonocardia,* its associated DxmA, *Mycobacterium* and its associated PrmA, were not detected in any of those sites (26). Therefore, while certain studies have also found a prevalence of Group V SDIMOs in dioxane-impacted sites, others have shown dioxane-degrading microbial communities that may utilize other SDIMO groups. These discrepancies may be due to environmental conditions such as redox potential (27).

## Conclusions

Support media from Lowry Landfill Bioreactor 1 showed stable and diverse microbial communities that contained potential dioxane degraders, namely *Pseudonocardia*. Lowry proteins that we were most confident encoded SDIMOs mostly belonged to Group V, with DxmA-like proteins being the most prevalent. Contigs containing DxmA-like Lowry proteins showed evidence of a range of horizontal gene transfer mechanisms. These may indicate a selection pressure in the bioreactor microbial communities to maintain Dxm. Additionally, other SDIMO groups were detected, showing the potential phylogenetic diversity of this one enzyme family in the bioreactor. Future studies may isolate dioxane-degrading species or simplified consortia from the bioreactor, test the dioxane degradation ability and mobility of the DxmA proteins from this work, and elucidate the overall microbial ecology of the Lowry Landfill BTS.

## ACKNOWLEDGEMENTS

We would like to thank the Lowry Trust for site access and permission to sample the bioreactors in the treatment plant of the Lowry Landfill Superfund Site. We thank Parsons Engineering for tours of the treatment plant, relaying site history including updates on remediation efforts, sampling assistance, and providing chemistry data from the treatment plant. We thank the many students enrolled in BIOL2021 at the University of Colorado Denver for help with 16S rRNA library preparation, and Daniel Frank, Jennifer Kofonow, and Cassandra Kotter for assistance with MiSeq sequencing.

This work was supported by the University of Colorado Denver 2020 Presidential Initiative on Urban and Place-Based Research. We thank Dr. Jan Mandel for access to the Alderaan cluster from the Center for Computational Mathematics at the University of Colorado Denver (supported by National Science Foundation award OAC-2019089).

## SUPPLEMENTAL TABLES

**Table S1.** Categories for the 39 SDIMOs described in K. L. Goff and L. A. Hug (15).

| **SDIMO category** | **Full description** | **Total** | **Sequence use in K. L. Goff and L. A. Hug (15)** |
| --- | --- | --- | --- |
| CDDP | Candidate dioxane degrading protein | 8 | Reference proteins from the literature with evidence of dioxane degradation that were searched for in genomes of known dioxane-degrading bacteria using BLASTp and in environmental metagenomes |
| OUT | Outgroup protein | 5 | Proteins from the literature with no evidence of dioxane degradation that were used as an anchor in phylogenetic trees |
| COMP | Composite protein | 17 | BLASTp-curated proteins from the genomes of known dioxane-degrading bacteria that shared either a high (>90%) or moderate (25-90%) percent identity with a CDDP |
| COMPOUT | Composite outgroup protein | 9 | BLASTp-curated proteins from the genomes of known dioxane-degrading bacteria that shared either a high (>90%) or moderate (25-90%) percent identity with a CDDP and were eventually presumed to not degrade dioxane based on phylogenetic placement |

**Table S2.** List of the 39 SDIMOs described in K. L. Goff and L. A. Hug (15).

|  | Abbreviation | Description | Accession | KO | Group | Length<br>(amino<br>acids) |
| --- | --- | --- | --- | --- | --- | --- |
| 1 | CDDP1 | *DxmA* [*Pseudonocardia dioxanivorans* CB1190] | WP_103383250.1 | K18223 | V | 545 |
| 2 | CDDP2 | *PrmA* [*Mycobacterium dioxanotrophicus* PH-06] | WP_087083743.1 | NA | VI | 512 |
| 3 | CDDP3 | *TmoA* [*Azoarcus* sp. DD4] | QDF97112.1 | K15760 | II | 501 |
| 4 | CDDP4 | *TomA3* [*Burkholderia cepacia* G4] | AAK07411.1 | K16242 | I | 519 |
| 5 | CDDP5 | *sMMO* [*Methylosinus trichosporium* OB3b] | ATQ70365.1 | K16157 | III | 526 |
| 6 | CDDP6 | *TmoA\** [*Pseudomonas mendocina* KR1] | AAA25999.1 | K15760 | II | 500 |
| 7 | CDDP7 | *TbuA1* [*Ralstonia pickettii* PKO1] | AAB09618.1 | K15760 | II | 501 |
| 8 | CDDP8 | *PrmA\** [*Rhodococcus* sp. RR1] | ADM83577.1 | K18223 | V | 439 |
| 9 | OUT1 | *alpha subunit-terminal oxygenase component* [*Pseudomonas* sp.] | AAA88459.1 | K16242 | I | 513 |
| 10 | OUT2 | *toluene, o-xylene monooxygenase oxygenase subunit* [*Pseudomonas* sp. OX1] | CAA06654.1 | K15760 | II | 498 |
| 11 | OUT3 | *Methane monooxygenase* [*Pseudonocardia dioxanivorans* CB1190] | AEA22892.1 | K18223 | V | 550 |
| 12 | OUT4 | *methane monooxygenase* [*Pseudonocardia acaciae*] | WP_156993170.1 | K18223 | V | 550 |
| 13 | OUT5 | *methane monooxygenase* [*Pseudonocardia asaccharolytica*] | WP_147201036.1 | K18223 | V | 550 |
| 14 | COMP1 | *butane monooxygenase alpha subunit* [*Brachymonas petroleovorans*] | AAR98534.1 | K16157 | III | 535 |

**Table S2.** List of the 39 SDIMOs described in K. L. Goff and L. A. Hug (15) (continued).
|  | Abbreviation | Description | Accession | KO | Group | Length<br>(amino<br>acids) |
| --- | --- | --- | --- | --- | --- | --- |
| 15 | COMP2 | *aromatic/alkene/methane*<br>*monooxygenase*<br>*hydroxylase/oxygenase*<br>*subunit alpha*<br>*[Methylococcus*<br>*capsulatus]* | WP_010960482.<br>1 | K16157 | III | 527 |
| 16 | COMP3 | *MULTISPECIES:*<br>*aromatic/alkene/methane*<br>*monooxygenase*<br>*hydroxylase/oxygenase*<br>*subunit alpha*<br>*[Methylosinus]* | WP_003609337.<br>1 | K16157 | III | 526 |
| 17 | COMP4 | *butane monooxygenase*<br>*hydroxylase BMOH alpha*<br>*subunit [Thauera*<br>*butanivorans]* | AAM19727.1 | K16157 | III | 530 |
| 18 | COMP5 | *aromatic/alkene/methane*<br>*monooxygenase*<br>*hydroxylase/oxygenase*<br>*subunit alpha*<br>*[Mycobacterium sp.*<br>*ENV421]* | WP_102810306.<br>1 | NA | VI | 513 |
| 19 | COMP6 | *MULTISPECIES:*<br>*aromatic/alkene/methane*<br>*monooxygenase*<br>*hydroxylase/oxygenase*<br>*subunit alpha*<br>*[Rhodococcus]* | WP_006947300.<br>1 | NA | VI | 512 |
| 20 | COMP7 | *soluble di-iron*<br>*monooxygenase alpha*<br>*subunit, partial*<br>*[Pseudonocardia sp. D17]* | BAU36819.1 | K18223 | V | 127 |
| 21 | COMP8 | *tetrahydrofuran*<br>*monooxygenase oxygenase*<br>*component alpha subunit*<br>*[Pseudonocardia sp.*<br>*ENV478]* | AEI99544.1 | K18223 | V | 545 |

**Table S2.** List of the 39 SDIMOs described in K. L. Goff and L. A. Hug (15) (continued).
|  | Abbreviation | Description | Accession | KO | Group | Length<br>(amino<br>acids) |
| --- | --- | --- | --- | --- | --- | --- |
| 22 | COMP9 | *methane monooxygenase*<br>[*Pseudonocardia* sp. N23] | WP_098956496.1 | K18223 | V | 545 |
| 23 | COMP10 | *alpha-subunit of multicomponent tetrahydrofuran monooxygenase*<br>[*Pseudonocardia tetrahydrofuranoxydans*] | CAC10506.1 | K18223 | V | 545 |
| 24 | COMP11 | *putative alkene monooxygenase alpha subunit*<br>[*Mycolicibacterium rhodesiae* JS60] | AAO48576.1 | K18223 | IV | 500 |
| 25 | COMP12 | *methane monooxygenase*<br>[*Pseudonocardia asaccharolytica*] | WP_028928894.1 | K18223 | V | 547 |
| 26 | COMP13 | *Propane 2-monooxygenase, hydroxylase component large subunit*<br>[*Rhodococcus jostii* RHA1] | Q0SJK9.1 | K18223 | V | 544 |
| 27 | COMP14 | *propane monooxygenase hydroxylase large subunit*<br>[*Gordonia* sp. TY-5] | BAD03956.2 | K18223 | V | 545 |
| 28 | COMP15 | *methane monooxygenase*<br>[*Mycobacterium goodii*] | WP_049747756.1 | K18223 | V | 542 |
| 29 | COMP16 | *methane monooxygenase*<br>[*Rhodococcus jostii*] | WP_011593714.1 | K18223 | V | 544 |
| 30 | COMP17 | *methane monooxygenase*<br>[*Mycolicibacterium smegmatis*] | WP_003893346.1 | K18223 | V | 542 |

**Table S2.** List of the 39 SDIMOs described in K. L. Goff and L. A. Hug (15) (continued).
|  | Abbreviation | Description | Accession | KO | Group | Length<br>(amino<br>acids) |
| --- | --- | --- | --- | --- | --- | --- |
| 31 | COMPOUT1 | *MULTISPECIES:*<br>*aromatic/alkene/methane*<br>*monooxygenase*<br>*hydroxylase/oxygenase*<br>*subunit alpha*<br>*[Acinetobacter]* | WP_033917438.1 | K16242 | I | 558 |
| 32 | COMPOUT2 | *MULTISPECIES:*<br>*aromatic/alkene/methane*<br>*monooxygenase*<br>*hydroxylase/oxygenase*<br>*subunit alpha*<br>*[Burkholderia cepacia*<br>*complex]* | WP_060217674.1 | K16242 | I | 513 |
| 33 | COMPOUT3 | *phenol 2-monooxygenase*<br>*[Ralstonia pickettii*<br>*DTP0602]* | AGW89993.1 | K16242 | I | 503 |
| 34 | COMPOUT4 | *Tbc1D monooxygenase*<br>*[Burkholderia cepacia]* | AAG40791.1 | K16242 | I | 514 |
| 35 | COMPOUT5 | *aromatic/alkene/methane*<br>*monooxygenase*<br>*hydroxylase/oxygenase*<br>*subunit alpha [Ralstonia*<br>*pickettii]* | WP_012430588.1 | K16242 | I | 514 |
| 36 | COMPOUT6 | *putative isoprene*<br>*monooxygenase alpha*<br>*subunit [Rhodococcus sp.*<br>*AD45]* | CAB55825.1 | K15760 | II | 514 |
| 37 | COMPOUT7 | *alkene monooxygenase*<br>*system oxygenase*<br>*component subunit alpha*<br>*[Xanthobacter*<br>*autotrophicus]* | WP_011992972.1 | K15760 | II | 497 |
| 38 | COMPOUT8 | *DMS oxygenase*<br>*component [Acinetobacter*<br>*sp.]* | BAA23333.1 | K16242 | I | 511 |
| 39 | COMPOUT9 | *Phenol 2-monooxygenase,*<br>*oxygenase component*<br>*DmpN [Pseudomonas sp.*<br>*CF600]* | P19732.1 | K16242 | I | 517 |

**Table S3.** Lowry Landfill Bioreactor 1 support media 16S rRNA sample (n=23) read counts and alpha diversity values.

|  | **Read Count** | **Observed** | **Shannon** | **Simpson** | **Faith PD** |
| --- | --- | --- | --- | --- | --- |
| 19-03-26-R1-N1-A | 66685 | 393 | 4.2 | 0.06 | 28 |
| 19-03-26-R1-N1-B | 134655 | 508 | 4.2 | 0.04 | 36 |
| 19-03-26-R1-N1-C | 100060 | 471 | 4.2 | 0.04 | 33 |
| 19-03-26-R1-N1-D | 13456 | 204 | 4.1 | 0.12 | 19 |
| 19-03-26-R1-N2-C | 48664 | 399 | 4.4 | 0.06 | 31 |
| 22-01-18-R1-N1-B | 12780 | 210 | 4.3 | 0.14 | 18 |
| 22-01-18-R1-N1-C | 5336 | 113 | 3.9 | 0.20 | 11 |
| 22-01-18-R1-N1-D | 16537 | 257 | 4.5 | 0.14 | 20 |
| 22-01-18-R1-N2-A | 23076 | 282 | 4.3 | 0.09 | 23 |
| 22-01-18-R1-N2-B | 8653 | 160 | 4.1 | 0.16 | 14 |
| 22-01-18-R1-N2-C | 7449 | 142 | 4.0 | 0.17 | 14 |
| 22-01-18-R1-N2-D | 26756 | 318 | 4.5 | 0.10 | 24 |
| 22-01-25-R1-N1-A | 28522 | 259 | 4.2 | 0.09 | 19 |
| 22-01-25-R1-N1-B | 13266 | 179 | 4.1 | 0.14 | 15 |
| 22-01-25-R1-N1-D | 10860 | 190 | 4.2 | 0.16 | 17 |
| 22-01-25-R1-N2-A | 10688 | 177 | 4.0 | 0.12 | 14 |
| 22-01-25-R1-N2-B | 7491 | 125 | 3.8 | 0.15 | 12 |
| 22-01-25-R1-N2-D | 19469 | 231 | 4.1 | 0.09 | 19 |
| 22-03-22-R1-N1-B | 14870 | 211 | 4.2 | 0.12 | 18 |
| 22-03-22-R1-N1-C | 6574 | 135 | 4.0 | 0.20 | 12 |
| 22-03-22-R1-N1-D | 10211 | 167 | 4.1 | 0.16 | 17 |
| 22-03-22-R1-N2-B | 10543 | 191 | 4.3 | 0.18 | 17 |
| 22-03-22-R1-N2-D | 13121 | 212 | 4.3 | 0.16 | 18 |

**Table S4.** Lowry Landfill Bioreactor 1 support media metagenomic shotgun sequencing sample (n=12) information and assembly statistics.

| **Sample Name** | **Date** | **Number of Contigs** | **Number of Contigs Greater Than or Equal to 1000 bp** | **N50** | **Largest Contig** |
| --- | --- | --- | --- | --- | --- |
| **S1** | 2019-03-26 | 1870969 | 313996 | 3044 | 1010632 |
| **S2** | 2019-03-26 | 2119006 | 371005 | 3408 | 1073174 |
| **S3** | 2019-03-26 | 1900365 | 313886 | 3386 | 1010632 |
| **S4** | 2022-01-18 | 2260283 | 347124 | 2999 | 1010632 |
| **S5** | 2022-01-18 | 2837071 | 429127 | 2622 | 1010632 |
| **S6** | 2022-01-18 | 2171795 | 344778 | 3058 | 767858 |
| **S7** | 2022-01-25 | 2399178 | 359486 | 2839 | 1010632 |
| **S8** | 2022-01-25 | 2272037 | 372938 | 2845 | 1010632 |
| **S9** | 2022-01-25 | 2438870 | 412304 | 3268 | 1190090 |
| **S10** | 2022-03-22 | 2527501 | 407399 | 3009 | 1010632 |
| **S11** | 2022-03-22 | 2652858 | 430838 | 3383 | 1010632 |
| **S12** | 2022-03-22 | 2626798 | 411297 | 3076 | 1010632 |

---

**Fig S1.**
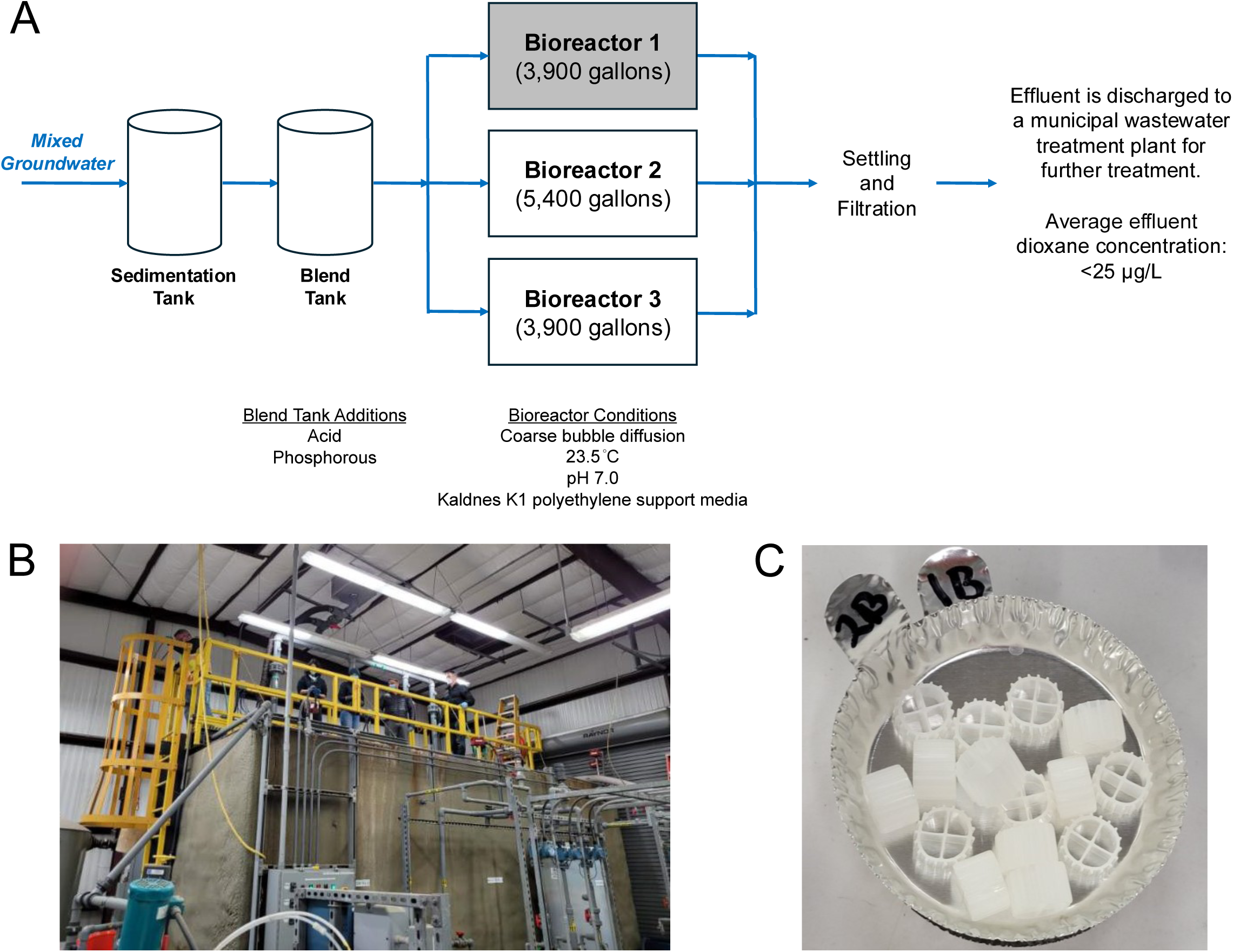
A schematic of the Lowry Landfill Biological Treatment System (BTS) adapted from L. Cordone et al. 2016 (12) (A). Chemistry data was collected from the Sedimentation Tank. Support media from Bioreactor 1 (shaded) were sequenced. Photo of the bioreactors (B). Photo of uninoculated Kaldnes K1 polyethylene support media (C).

**Fig S2.**
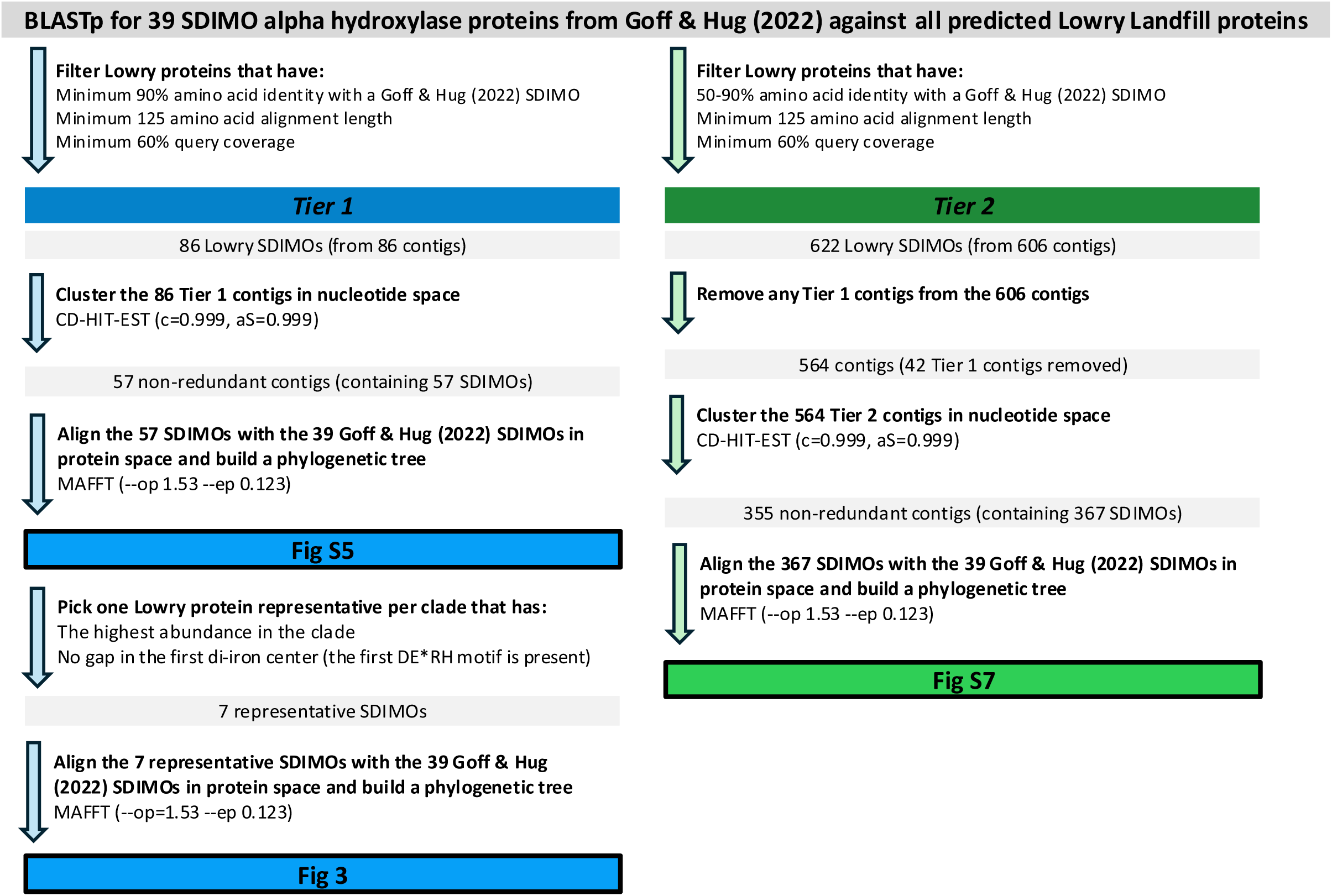
Flowchart of the tiered approach to rank SDIMO proteins curated from Lowry Landfill samples.

**Fig S3.**
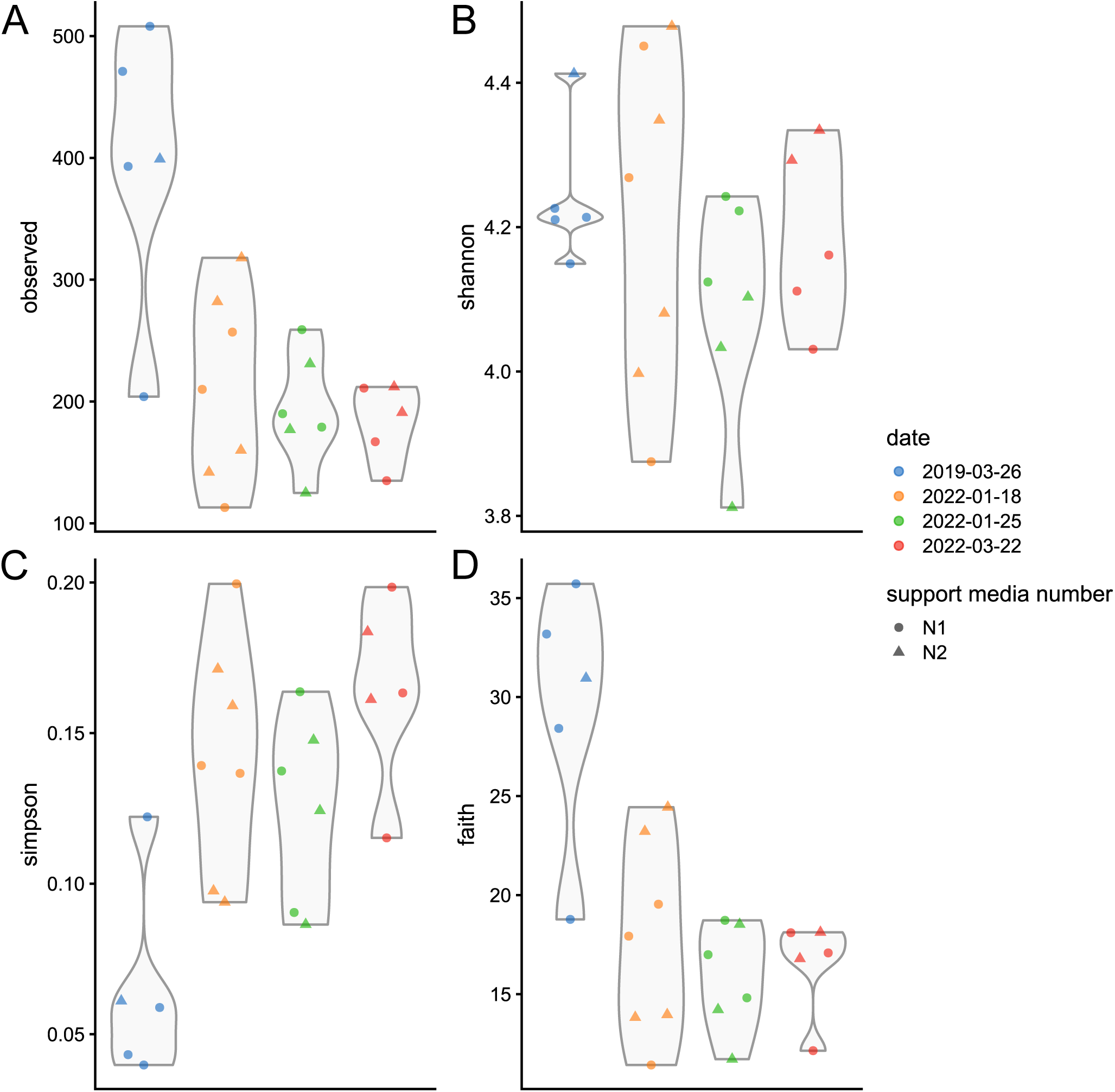
Alpha diversity analyses for Lowry Landfill Bioreactor 1 support media (n=23). The analyses performed included Observed Richness (A), Shannon (B), Simpson (C), and Faith PD (D) metrics.

**Fig S4.**
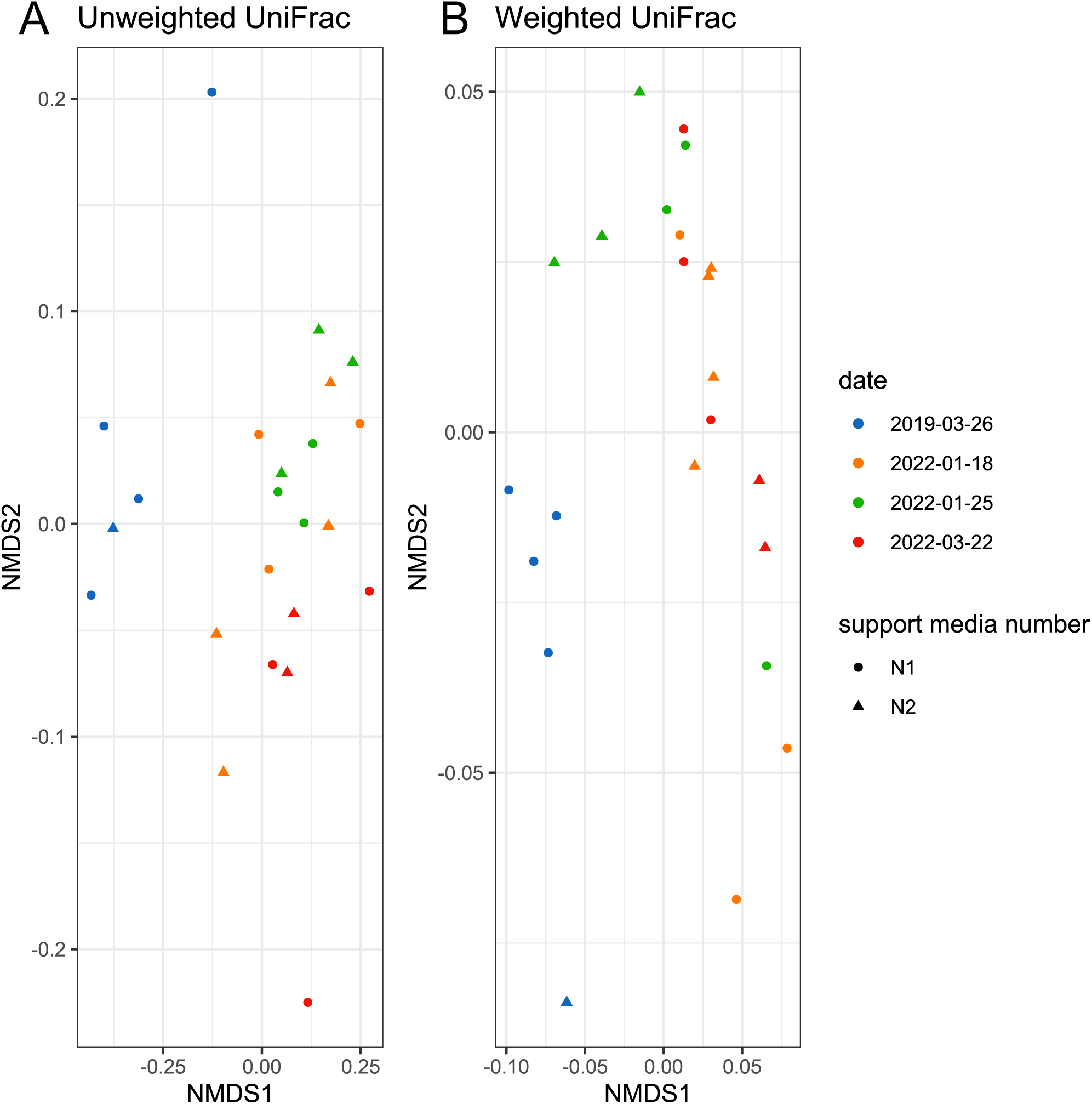
Beta diversity analyses for Lowry Landfill Bioreactor 1 support media (n=23). Unweighted (excluding taxon abundances, A) and weighted (including taxon abundances, B) non-metric multidimensional scaling (NMDS) analyses were based on UniFrac distances.

**Fig S5.**
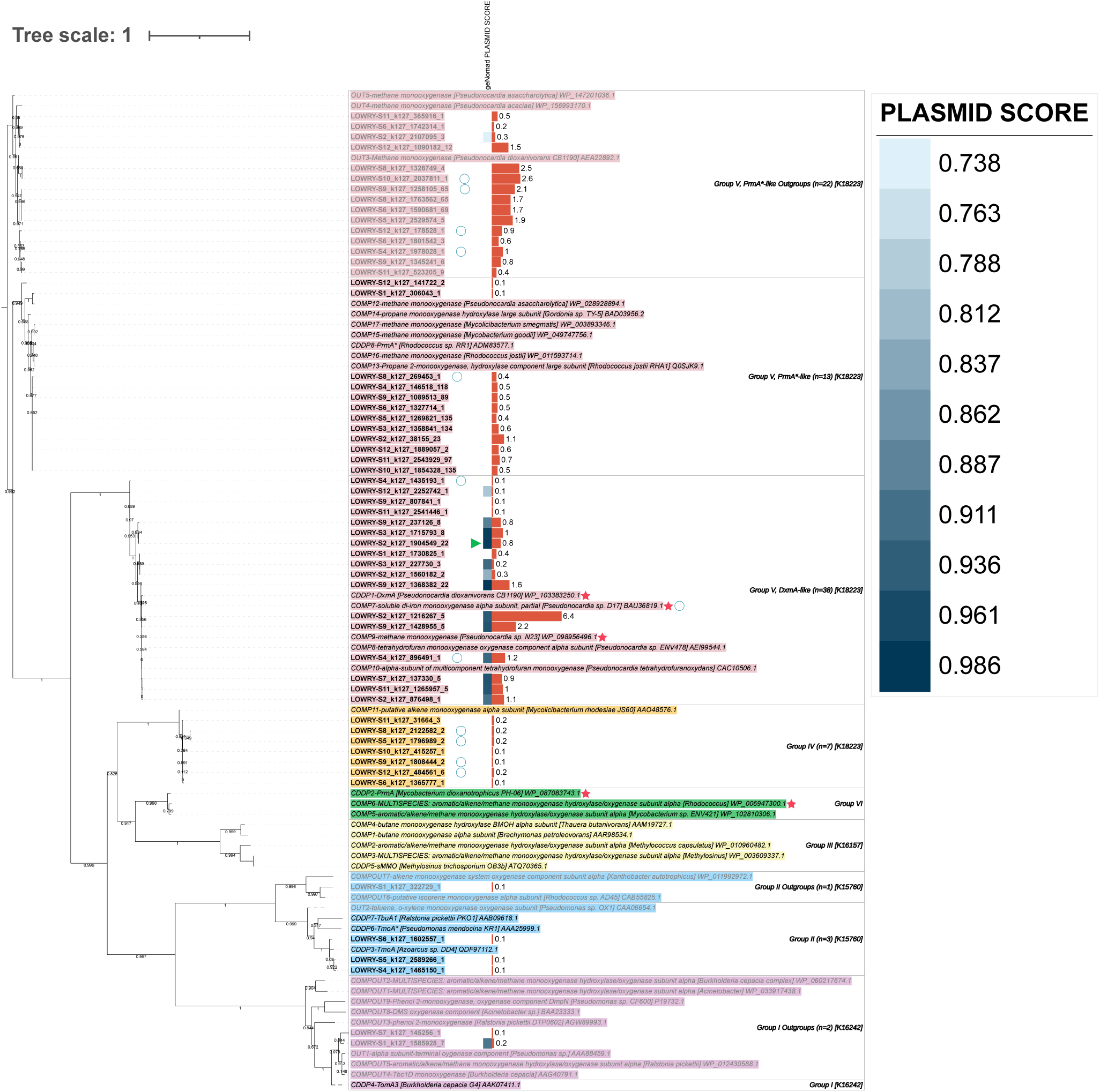
Protein phylogenetic tree of sequences recovered from Lowry Landfill Bioreactor 1 support media within 57 non-redundant contigs and 39 candidate SDIMOs described by K. L. Goff and L. A. Hug (15). Protein abundances were estimated using coverages (RPKM) of the 86 original Lowry contigs containing SDIMOs and displayed in bar charts for each sequence. Sequences containing an inverted terminal repeat according to geNomad are marked with a green triangle. Plasmid scores above the geNomad default threshold are displayed in the heatmap. See Fig. 3 legend for more details on symbols and labels.

**Fig S6.**
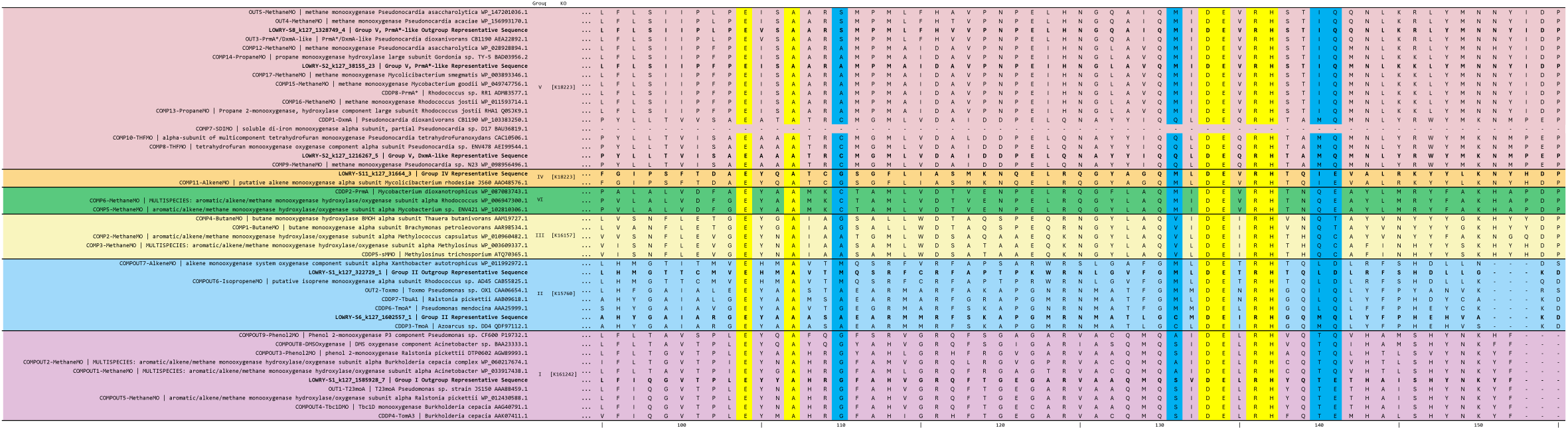
Amino acid alignment of the first region of the di-iron center of the alpha hydroxylase subunit of seven Lowry Landfill representative protein sequences and 39 candidate SDIMOs described by K. L. Goff and L. A. Hug (15) (positions 96-155). Residues highlighted in yellow showed high conservation. Residues highlighted in blue indicate hydrophobic amino acids used to classify sequences into SDIMO Groups I-VI.

**Fig S7.**
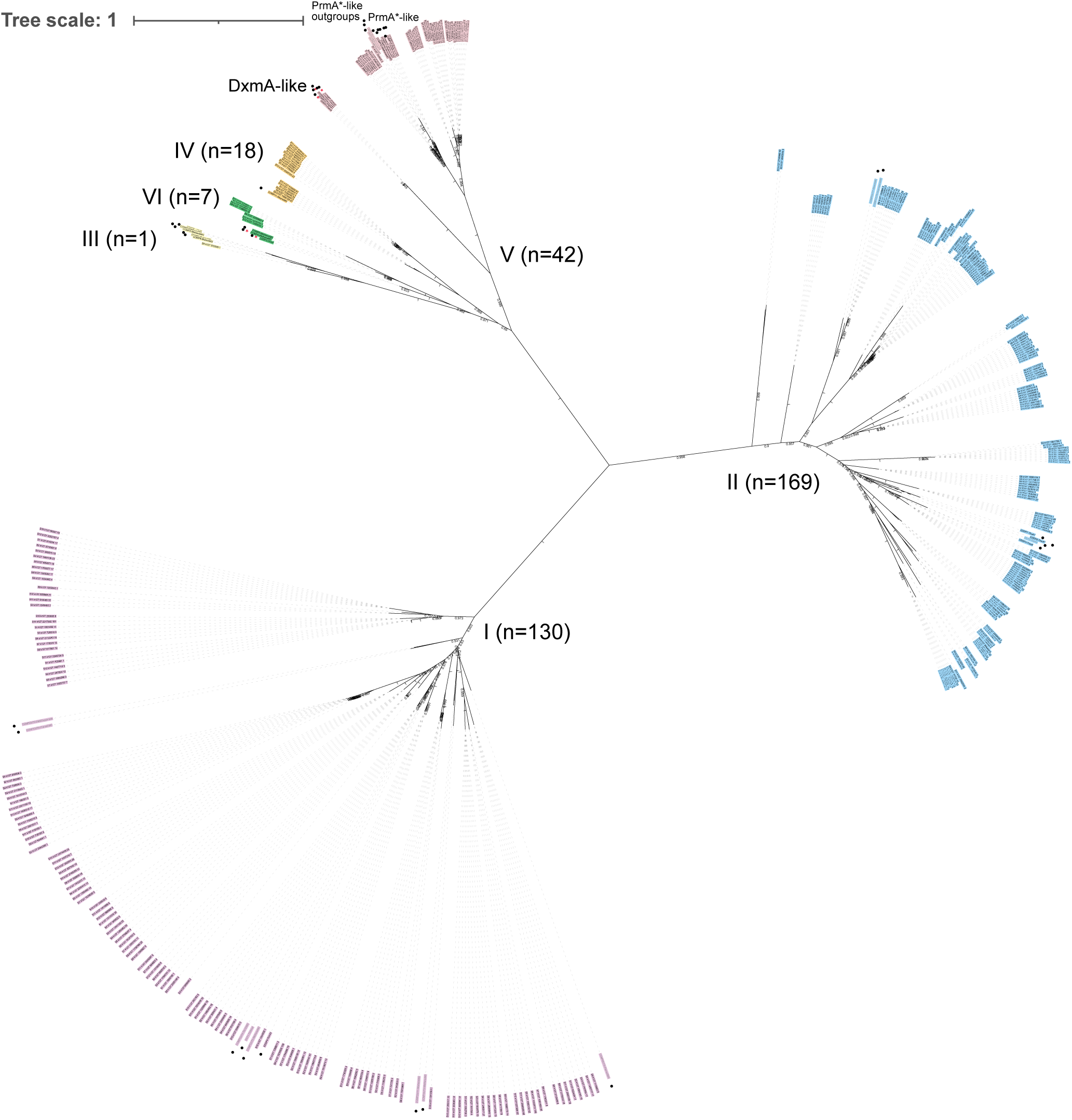
Unrooted protein phylogenetic tree of 367 Tier 2 potential SDIMO sequences recovered from Lowry Landfill Bioreactor 1 support media and 39 candidate SDIMOs described by K. L. Goff and L. A. Hug (15). The 367 potential SDIMO sequences were recovered from a set of 355 non-redundant contigs. Proteins from K. L. Goff and L. A. Hug (15) that are presumed not to degrade dioxane due to monophyletic clading with known outgroups are written in gray text. The branches for OUT2 and CDDP4 were dashed as these showed monophyletic clading that was unexpected for their description in K. L. Goff and L. A. Hug (15). Red stars indicate sequences that have shown direct metabolism of dioxane in the literature. Circles indicate sequences from K. L. Goff and L. A. Hug (15) whereas unmarked sequences were recovered from Lowry Landfill. Lowry sequences are colored by predicted SDIMO group based on clading patterns with literature sequences.

**Fig S8.**
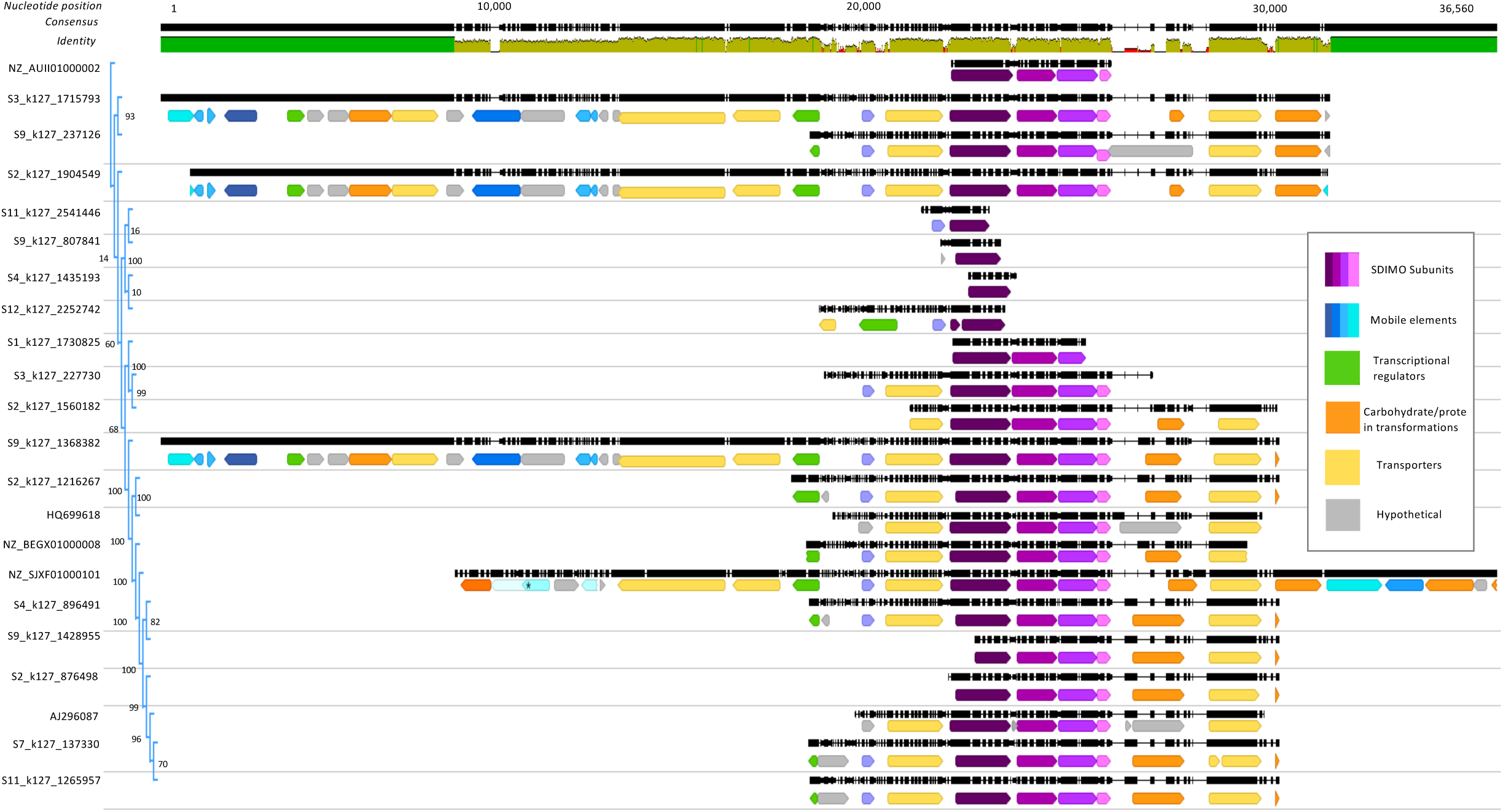
Alignment and phylogeny of the Lowry DxmA-like CDDP containing contigs, along with representative contigs from K. L. Goff and L. A. Hug (15). Predicted proteins and annotation categories are color coded by function (see inset). On the identity plot, green indicates 100% identity, while greenish brown indicates at least 30% and below 100%, and red indicates below 30%. Bootstrap values are denoted at the nodes of the phylogenetic tree.

## REFERENCES

1. EPA. 2017. Technical Fact Sheet – 1,4-Dioxane (EPA-505-F-17-011).

2. EPA. 2006. Treatment Technologies for 1,4-Dioxane: Fundamentals and Field Applications.

3. Pollitt KJG, Kim JH, Peccia J, Elimelech M, Zhang YW, Charkoftaki G, Hodges B, Zucker I, Huang H, Deziel NC, Murphy K, Ishii M, Johnson CH, Boissevain A, O’Keefe E, Anastas PT, Orlicky D, Thompson DC, Vasiliou V. 2019. 1,4-Dioxane as an emerging water contaminant: State of the science and evaluation of research needs. Science of the Total Environment 690:853–866.

4. Mohr TKGD, W.; Hatton, J.; Anderson, J. 2020. Environmental Investigation and Remediation: 1,4-Dioxane and other Solvent Stabilizers, Second ed doi:https://doi-org.aurarialibrary.idm.oclc.org/10.1201/9780429401428, Boca Raton.

5. McElroy AC, Hyman MR, Knappe DRU. 2019. 1,4-Dioxane in drinking water: emerging for 40 years and still unregulated. Current Opinion in Environmental Science & Health 7:117–125.

6. EPA. 2018. Problem Formulation of the Risk Evaluation for 1,4-Dioxane.

7. Doherty AC, Lee CS, Meng QY, Sakano Y, Noble AE, Grant KA, Esposito A, Gobler CJ, Venkatesan AK. 2023. Contribution of household and personal care products to 1,4-dioxane contamination of drinking water. Current Opinion in Environmental Science & Health 31.

8. Zenker MJ, Borden RC, Barlaz MA. 2003. Occurrence and treatment of 1,4-dioxane in aqueous environments. Environmental Engineering Science 20:423–432.

9. Ramos P, Kwok IY, Ngo J, Zgonc D, Miao Y, Pornwongthong P, Blotevogel J, Mahendra S. 2022. A, B, Cs of 1,4-dioxane removal from water: Adsorption, biodegradation, and catalysis. Current Opinion in Environmental Science & Health 29.

10. Dawson D, Fisher H, Noble AE, Meng QY, Doherty AC, Sakano Y, Vallero D, Tornero-Velez R, Hubal EAC. 2022. Assessment of Non-Occupational 1,4-Dioxane Exposure Pathways from Drinking Water and Product Use. Environmental Science & Technology 56:5266–5275.

11. Kim CG, Seo HJ, Lee BR. 2006. Decomposition of 1,4-dioxane by advanced oxidation and biochemical process. Journal of Environmental Science and Health Part a-Toxic/Hazardous Substances & Environmental Engineering 41:599–611.

12. Cordone L, Carlson C, Plaehn W, Shangraw T, Wilmoth D. 2016. Case Study and Retrospective: Aerobic Fixed Film Biological Treatment Process for 1,4-Dioxane at the Lowry Landfill Superfund Site. Remediation Journal 27:159–172.

13. Zhang S, Gedalanga PB, Mahendra S. 2017. Advances in bioremediation of 1,4-dioxane-contaminated waters. Journal of Environmental Management 204:765–774.

14. Maier RM. 2019. Biological Processes Affecting Contaminants Transport and Fate, p 131–146. *In* Mark L. Brusseau ILP, Charles P. Gerba (ed), Environmental and Pollution Science, 3 ed 10.1016/B978-0-12-814719-1.00009-4. Academic Press.

15. Goff KL, Hug LA. 2022. Environmental Potential for Microbial 1,4-Dioxane Degradation Is Sparse despite Mobile Elements Playing a Role in Trait Distribution. Applied and Environmental Microbiology 88:e02091–21.

16. Tang YY, Mao XW. 2023. Recent Advances in 1,4-Dioxane Removal Technologies for Water and Wastewater Treatment. Water 15.

17. Parales RE, Adamus JE, White N, May HD. 1994. Degradation of 1,4-Dioxane by an Actinomycete in Pure Culture. Applied and Environmental Microbiology 60:4527–4530.

18. Mahendra S, Alvarez-Cohen L. 2006. Kinetics of 1,4-dioxane biodegradation by monooxygenase-expressing bacteria. Environmental Science & Technology 40:5435–5442.

19. Chen RH, Miao Y, Liu Y, Zhang L, Zhong M, Adams JM, Dong YH, Mahendra S. 2021. Identification of novel 1,4-dioxane degraders and related genes from activated sludge by taxonomic and functional gene sequence analysis. Journal of Hazardous Materials 412.

20. Dai CH, Wu H, Wang XJ, Zhao KK, Lu ZM. 2022. Network and meta-omics reveal the cooperation patterns and mechanisms in an efficient 1,4-dioxane-degrading microbial consortium. Chemosphere 301.

21. Hassan R, Kriaa K, Wahaballa AM, Elsayed M, Mahmoud M, Nasr M, Tawfik A. 2024. Performance assessment of up-flow anaerobic multi-staged reactor followed by auto-aerated immobilized biomass unit for treating polyester wastewater, with biogas production. Applied Water Science 14.

22. Li F, Deng DY, Wadden A, Parvis P, Cutt D, Li MY. 2023. Effective removal of trace 1,4-dioxane by biological treatments augmented with propanotrophic single culture versus synthetic consortium. Journal of Hazardous Materials Advances 9.

23. Holmes AJ. 2009. The Diversity of Soluble Di-iron Monooxygenases with Bioremediation Applications. Advances in Applied Bioremediation 17:91–102.

24. Sadeghi V, Mora R, Jacob P, Chiang S-YD. 2016. CharacterizingIn SituMethane-Enhanced Biostimulation Potential for 1,4-Dioxane Biodegradation in Groundwater. Remediation Journal 27:115–132.

25. Li MY, Mathieu J, Yang Y, Fiorenza S, Deng Y, He ZL, Zhou JZ, Alvarez PJJ. 2013. Widespread Distribution of Soluble Di-Iron Monooxygenase (SDIMO) Genes in Arctic Groundwater Impacted by 1,4-Dioxane. Environmental Science & Technology 47:9950–9958.

26. Dang H, Kanitkar YH, Stedtfeld RD, Hatzinger PB, Hashsham SA, Cupples AM. 2018. Abundance of Chlorinated Solvent and 1,4-Dioxane Degrading Microorganisms at Five Chlorinated Solvent Contaminated Sites Determined via Shotgun Sequencing. Environmental Science &amp; Technology 52:13914–13924.

27. Ramalingam V, Cupples AM. 2020. Enrichment of novel Actinomycetales and the detection of monooxygenases during aerobic 1,4-dioxane biodegradation with uncontaminated and contaminated inocula. Applied Microbiology and Biotechnology 104:2255–2269.

28. EPA. n.d. Superfund Site: LOWRY LANDFILL UNINCORPORATED ARAPAHOE COUNTY, CO. https://cumulis.epa.gov/supercpad/SiteProfiles/index.cfm?fuseaction=second.Cleanup&id=0800186#content. Accessed October 27, 2023.

29. N/A. n.d. Lowry Landfill. https://lowrylandfill.com/. Accessed October 27, 2023.

30. EPA. 2007. LOWRY LANDFILL SUPERFUND SITE: THIRD EXPLANATION OF SIGNIFICANT DIFFERENCES. Environmental Protection Agency,

31. Roane TM. 2025. Subject: [EXTERNAL] Lowry paper (Lowry communication).

32. EPA. 2017. FOURTH FIVE-YEAR REVIEW REPORT FOR LOWRY LANDFILL SUPERFUND SITE. Environmental Protection Agency, Denver, CO.

33. EPA. 1996. METHOD 8260B. https://19january2017snapshot.epa.gov/sites/production/files/2015-12/documents/8260b.pdf.

34. Parada AE, Needham DM, Fuhrman JA. 2016. Every base matters: assessing small subunit rRNA primers for marine microbiomes with mock communities, time series and global field samples. Environmental Microbiology 18:1403–1414.

35. Apprill A, McNally S, Parsons R, Weber L. 2015. Minor revision to V4 region SSU rRNA 806R gene primer greatly increases detection of SAR11 bacterioplankton. Aquatic Microbial Ecology 75:129–137.

36. Kozich JJ, Westcott SL, Baxter NT, Highlander SK, Schloss PD. 2013. Development of a Dual-Index Sequencing Strategy and Curation Pipeline for Analyzing Amplicon Sequence Data on the MiSeq Illumina Sequencing Platform. Applied and Environmental Microbiology 79:5112–5120.

37. Bokulich NA, Kaehler BD, Rideout JR, Dillon M, Bolyen E, Knight R, Huttley GA, Caporaso JG. 2018. Optimizing taxonomic classification of marker-gene amplicon sequences with QIIME 2′s q2-feature-classifier plugin. Microbiome 6.

38. Robeson MS, O’Rourke, D.R., Kaehler, B.D., Ziemski, M., Dillon, M.R., Foster, J.T., Bokulich, N.A. 2020. RESCRIPt: Reproducible sequence taxonomy reference database management for the masses. 10.1101/2020.10.05.326504.

39. Ernst F., Shetty, S., Borman, T., Lahti, L. 2023. mia: Microbiome analysis, v1.8.0. https://bioconductor.org/packages/mia.

40. Morgan M., Wang, J., Obenchain, V., Lang, M., Thompson, R., Turaga, N. 2023. BiocParallel: Bioconductor facilities for parallel evaluation, v1.34.2. https://bioconductor.org/packages/BiocParallel.

41. Wickham H. 2022. stringr: Simple, Consistent Wrappers for Common String Operations, v1.5.0. https://stringr.tidyverse.org.

42. Pagès H, Lawrence, M., Aboyoun, P. 2023. S4Vectors: Foundation of vector-like and list-like containers in Bioconductor, v0.38.1. https://bioconductor.org/packages/S4Vectors.

43. Huber W, Carey VJ, Gentleman R, Anders S, Carlson M, Carvalho BS, Bravo HC, Davis S, Gatto L, Girke T, Gottardo R, Hahne F, Hansen KD, Irizarry RA, Lawrence M, Love MI, MacDonald J, Obenchain V, Oles AK, Pagès H, Reyes A, Shannon P, Smyth GK, Tenenbaum D, Waldron L, Morgan M. 2015. Orchestrating high-throughput genomic analysis with Bioconductor. Nature Methods 12:115–121.

44. McCarthy DJ, Campbell KR, Lun ATL, Wills QF. 2017. Scater: pre-processing, quality control, normalization and visualization of single-cell RNA-seq data in R. Bioinformatics 33:1179–1186.

45. Pedersen T. 2023. patchwork: The Composer of Plots, v1.1.2. https://patchwork.data-imaginist.com, https://github.com/thomasp85/patchwork.

46. McMurdie PJ, Holmes S. 2013. phyloseq: An R Package for Reproducible Interactive Analysis and Graphics of Microbiome Census Data. Plos One 8.

47. Wickham H. 2016. ggplot2: Elegant Graphics for Data Analysis, Springer International Publishing, https://ggplot2.tidyverse.org/.

48. Pedersen T. 2022. ggraph: An Implementation of Grammar of Graphics for Graphs and Networks, v2.1.0. https://ggraph.data-imaginist.com, https://github.com/thomasp85/ggraph.

49. Bushnell B, Rood J, Singer E. 2017. BBMerge - Accurate paired shotgun read merging via overlap. Plos One 12.

50. Andrews S. 2010. FastQC: A Quality Control Tool for High Throughput Sequence Data. http://www.bioinformatics.babraham.ac.uk/projects/fastqc/. Accessed November 25, 2024.

51. Ewels P, Magnusson M, Lundin S, Käller M. 2016. MultiQC: summarize analysis results for multiple tools and samples in a single report. Bioinformatics 32:3047–3048.

52. Li DH, Liu CM, Luo RB, Sadakane K, Lam TW. 2015. MEGAHIT: an ultra-fast single-node solution for large and complex metagenomics assembly via succinct graph. Bioinformatics 31:1674–1676.

53. Gurevich A, Saveliev V, Vyahhi N, Tesler G. 2013. QUAST: quality assessment tool for genome assemblies. Bioinformatics 29:1072–1075.

54. Li H, Handsaker B, Wysoker A, Fennell T, Ruan J, Homer N, Marth G, Abecasis G, Durbin R, Proc GPD. 2009. The Sequence Alignment/Map format and SAMtools. Bioinformatics 25:2078–2079.

55. Hyatt D, Chen GL, LoCascio PF, Land ML, Larimer FW, Hauser LJ. 2010. Prodigal: prokaryotic gene recognition and translation initiation site identification. Bmc Bioinformatics 11.

56. Camacho C, Coulouris G, Avagyan V, Ma N, Papadopoulos J, Bealer K, Madden TL. 2009. BLAST plus : architecture and applications. Bmc Bioinformatics 10.

57. Kanehisa M, Sato Y, Morishima K. 2016. BlastKOALA and GhostKOALA: KEGG Tools for Functional Characterization of Genome and Metagenome Sequences. Journal of Molecular Biology 428:726–731.

58. Inoue D, Tsunoda T, Sawada K, Yamamoto N, Saito Y, Sei K, Ike M. 2016. 1,4-Dioxane degradation potential of members of the genera Pseudonocardia and Rhodococcus. Biodegradation 27:277–286.

59. Katoh K, Standley DM. 2013. MAFFT Multiple Sequence Alignment Software Version 7: Improvements in Performance and Usability. Molecular Biology and Evolution 30:772–780.

60. Leahy JG, Batchelor PJ, Morcomb SM. 2003. Evolution of the soluble diiron monooxygenases. Fems Microbiology Reviews 27:449–479.

61. Price MN, Dehal PS, Arkin AP. 2009. FastTree: Computing Large Minimum Evolution Trees with Profiles instead of a Distance Matrix. Molecular Biology and Evolution 26:1641–1650.

62. Letunic I, Bork P. 2021. Interactive Tree Of Life (iTOL) v5: an online tool for phylogenetic tree display and annotation. Nucleic Acids Research 49:W293–W296.

63. Aroney STN, Newell, R. J. P., Nissen, J., Camargo, A. P., Tyson, G. W., & Woodcroft, B. J. 2024. CoverM: Read coverage calculator for metagenomics, on Zenodo. 10.5281/zenodo.10531253. Accessed November 25, 2024.

64. Li WZ, Godzik A. 2006. Cd-hit: a fast program for clustering and comparing large sets of protein or nucleotide sequences. Bioinformatics 22:1658–1659.

65. Fu LM, Niu BF, Zhu ZW, Wu ST, Li WZ. 2012. CD-HIT: accelerated for clustering the next-generation sequencing data. Bioinformatics 28:3150–3152.

66. He Y, Mathieu J, Yang Y, Yu PF, da Silva MLB, Alvarez PJJ. 2017. 1,4-Dioxane Biodegradation by Mycobacterium dioxanotrophicus PH-06 Is Associated with a Group-6 Soluble Di-Iron Monooxygenase. Environmental Science & Technology Letters 4:494–499.

67. Li MY, Mathieu J, Liu YY, Van Orden ET, Yang Y, Fiorenza S, Alvarez PJJ. 2014. The Abundance of Tetrahydrofuran/Dioxane Monooxygenase Genes (thmA/dxmA) and 1,4-Dioxane Degradation Activity Are Significantly Correlated at Various Impacted Aquifers. Environmental Science & Technology Letters 1:122–127.

68. Arkin AP, Cottingham RW, Henry CS, Harris NL, Stevens RL, Maslov S, Dehal P, Ware D, Perez F, Canon S, Sneddon MW, Henderson ML, Riehl WJ, Murphy-Olson D, Chan SY, Kamimura RT, Kumari S, Drake MM, Brettin TS, Glass EM, Chivian D, Gunter D, Weston DJ, Allen BH, Baumohl J, Best AA, Bowen B, Brenner SE, Bun CC, Chandonia JM, Chia JM, Colasanti R, Conrad N, Davis JJ, Davison BH, DeJongh M, Devoid S, Dietrich E, Dubchak I, Edirisinghe JN, Fang G, Faria JP, Frybarger PM, Gerlach W, Gerstein M, Greiner A, Gurtowski J, Haun HL, He F, Jain R, et al. 2018. KBase: The United States Department of Energy Systems Biology Knowledgebase. Nature Biotechnology 36:566–569.

69. Shaffer M, Borton MA, Bolduc B, Faria JP, Flynn RM, Ghadermazi P, Edirisinghe JN, Wood-Charlson EM, Miller CS, Chan SHJ, Sullivan MB, Henry CS, Wrighton KC. 2023. kb_DRAM: annotation and metabolic profiling of genomes with DRAM in KBase. Bioinformatics 39.

70. Brettin T, Davis JJ, Disz T, Edwards RA, Gerdes S, Olsen GJ, Olson R, Overbeek R, Parrello B, Pusch GD, Shukla M, Thomason JA, 3rd, Stevens R, Vonstein V, Wattam AR, Xia F. 2015. RASTtk: a modular and extensible implementation of the RAST algorithm for building custom annotation pipelines and annotating batches of genomes. Sci Rep 5:8365.

71. Shaffer M, Borton MA, McGivern BB, Zayed AA, La Rosa SL, Solden LM, Liu P, Narrowe AB, Rodriguez-Ramos J, Bolduc B, Gazitua MC, Daly RA, Smith GJ, Vik DR, Pope PB, Sullivan MB, Roux S, Wrighton KC. 2020. DRAM for distilling microbial metabolism to automate the curation of microbiome function. Nucleic Acids Res 48:8883–8900.

72. Aziz RK, Bartels D, Best AA, DeJongh M, Disz T, Edwards RA, Formsma K, Gerdes S, Glass EM, Kubal M, Meyer F, Olsen GJ, Olson R, Osterman AL, Overbeek RA, McNeil LK, Paarmann D, Paczian T, Parrello B, Pusch GD, Reich C, Stevens R, Vassieva O, Vonstein V, Wilke A, Zagnitko O. 2008. The RAST Server: rapid annotations using subsystems technology. BMC Genomics 9:75.

73. Kanehisa M, Goto S. 2000. KEGG: kyoto encyclopedia of genes and genomes. Nucleic Acids Res 28:27–30.

74. Mistry J, Chuguransky S, Williams L, Qureshi M, Salazar GA, Sonnhammer ELL, Tosatto SCE, Paladin L, Raj S, Richardson LJ, Finn RD, Bateman A. 2021. Pfam: The protein families database in 2021. Nucleic Acids Res 49:D412–D419.

75. Camargo AP, Roux S, Schulz F, Babinski M, Xu Y, Hu B, Chain PSG, Nayfach S, Kyrpides NC. 2023. Identification of mobile genetic elements with geNomad. Nature Biotechnology 10.1038/s41587-023-01953-y.

76. Ross K, Varani AM, Snesrud E, Huang H, Alvarenga DO, Zhang J, Wu C, McGann P, Chandler M. 2021. TnCentral: a Prokaryotic Transposable Element Database and Web Portal for Transposon Analysis. mBio 12:e0206021.

77. Guan J, Chen Y, Goh YX, Wang M, Tai C, Deng Z, Song J, Ou HY. 2024. TADB 3.0: an updated database of bacterial toxin-antitoxin loci and associated mobile genetic elements. Nucleic Acids Res 52:D784–D790.

78. Crescente JM, Zavallo D, Helguera M, Vanzetti LS. 2018. MITE Tracker: an accurate approach to identify miniature inverted-repeat transposable elements in large genomes. BMC Bioinformatics 19:348.

79. Stamatakis A. 2014. RAxML version 8: a tool for phylogenetic analysis and post-analysis of large phylogenies. Bioinformatics 30:1312–1313.

80. Deng DY, Li F, Li MY. 2018. A Novel Propane Monooxygenase Initiating Degradation of 1,4-Dioxane by Mycobacterium dioxanotrophicus PH-06. Environmental Science & Technology Letters 5:86–91.

81. Grostern A, Sales CM, Zhuang WQ, Erbilgin O, Alvarez-Cohen L. 2012. Glyoxylate Metabolism Is a Key Feature of the Metabolic Degradation of 1,4-Dioxane by Pseudonocardia dioxanivorans Strain CB1190. Applied and Environmental Microbiology 78:3298–3308.

82. Yamamoto N, Saito Y, Inoue D, Sei K, Ike M. 2018. Characterization of newly isolated sp N23 with high 1,4-dioxane-degrading ability. Journal of Bioscience and Bioengineering 125:552–558.

83. Vainberg S, McClay K, Masuda H, Root D, Condee C, Zylstra GJ, Steffan RJ. 2006. Biodegradation of ether pollutants by sp strain ENV478. Applied and Environmental Microbiology 72:5218–5224.

84. Hand S, Wang BX, Chu KH. 2015. Biodegradation of 1,4-dioxane: Effects of enzyme inducers and trichloroethylene. Science of the Total Environment 520:154–159.

85. Deng DY, Pham DN, Li F, Li MY. 2020. Discovery of an Inducible Toluene Monooxygenase That Cooxidizes 1,4-Dioxane and 1,1-Dichloroethylene in Propanotrophic Azoarcus sp. Strain DD4. Applied and Environmental Microbiology 86.

86. Chauvaux S, Chevalier F, Dantec CL, Fayolle F, Miras I, Kunst F, Beguin P. 2001. Cloning of a Genetically Unstable Cytochrome P-450 Gene Cluster Involved in Degradation of the Pollutant Ethyl tert-Butyl Ether by Rhodococcus ruber. Journal of Bacteriology 183:6551–6557.

87. Fraikin N, Goormaghtigh F, Van Melderen L. 2020. Type II Toxin-Antitoxin Systems: Evolution and Revolutions. J Bacteriol 202.

88. Jurenas D, Fraikin N, Goormaghtigh F, Van Melderen L. 2022. Biology and evolution of bacterial toxin-antitoxin systems. Nat Rev Microbiol 20:335–350.

89. Groth AC, Calos MP. 2004. Phage integrases: biology and applications. J Mol Biol 335:667–78.

90. Nadal-Molero F, Rosselli R, Garcia-Juan S, Campos-Lopez A, Martin-Cuadrado AB. 2024. Unveiling host-parasite relationships through conserved MITEs in prokaryote and viral genomes. Nucleic Acids Research 52:13094–13109.

91. Minnick MF. 2024. Functional Roles and Genomic Impact of Miniature Inverted-Repeat Transposable Elements (MITEs) in Prokaryotes. Genes (Basel) 15.

92. Gedalanga P, Madison A, Miao Y, Richards T, Hatton J, Diguiseppi WH, Wilson J, Mahendra S. 2016. A Multiple Lines of Evidence Framework to Evaluate Intrinsic Biodegradation of 1,4-Dioxane. Remediation Journal 27:93–114.

93. Tinberg CE, Lippard SJ. 2011. Dioxygen Activation in Soluble Methane Monooxygenase. Accounts of Chemical Research 44:280–288.

94. Li F, Deng DY, Li MY. 2020. Distinct Catalytic Behaviors between Two 1,4-Dioxane-Degrading Monooxygenases: Kinetics, Inhibition, and Substrate Range. Environmental Science & Technology 54:1898–1908.

95. He Y, Mathieu J, da Silva MLB, Li MY, Alvarez PJJ. 2018. 1,4-Dioxane-degrading consortia can be enriched from uncontaminated soils: prevalence of Mycobacterium and soluble di-iron monooxygenase genes. Microbial Biotechnology 11:189–198.

